# Chemical N-degrons activate p62-mediated mitophagy to alleviate mitochondrial neuropathies

**DOI:** 10.64898/2026.02.19.706908

**Authors:** Soon Chul Kwon, Byeong-Seong Kim, Hyeyoon Kim, Dae Eun Kang, Gee Eun Lee, Eui Jung Jung, Min Ju Lee, Yeon Sung Son, Da-ha Park, Daniel Youngjae Park, Jihoon Lee, Eun Hye Cho, Su Bin Kim, Ah Jung Heo, Young Ho Suh, Woo-Dong Jang, Dohyun Han, Chang Hoon Ji, Jee-Yin Ahn, Yong Tae Kwon

**Affiliations:** Department of Biomedical Sciences, College of Medicine, Seoul National University, Seoul 03080, Republic of Korea; Cellular Degradation Biology Center, College of Medicine, Seoul National University, Seoul 03080, Republic of Korea; Department of Molecular Cell Biology, Sungkyunkwan University School of Medicine, Suwon 16419, Republic of Korea; Department of Transdisciplinary Medicine, Seoul National University Hospital, Seoul 03082, Republic of Korea; College of Pharmacy, Sookmyung Women’s University, Seoul 04310, Republic of Korea; Department of Chemistry, Yonsei University, Seoul 03722, Republic of Korea; Division of Molecular Medicine, Department of Medicine, Columbia University Irving Medical Center, NY 10032, USA; Ischemic/Hypoxic Disease Institute, Medical Research Center, Seoul National University, Seoul 03080, Republic of Korea; Neuroscience Research Institute, Medical Research Center, Seoul National University, Seoul 03080, Republic of Korea; Institute of Biochemistry II, Faculty of Medicine, Goethe University Frankfurt, Frankfurt am Main 60590, Germany; AUTOTAC Bio Inc., Seoul 08501, Republic of Korea; Samsung Biomedical Research Institute, Samsung Medical Center, Seoul 06351, Republic of Korea; Convergence Dementia Research Center, Medical Research Center, Seoul National University, Seoul 03080, Republic of Korea; National Research Laboratory for Convergence Degradation Biology, Korea University, Seoul 02841, Republic of Korea

## Abstract

Pharmacological activation of mitophagy offers a promising strategy to eliminate dysfunctional mitochondria. We previously identified the autophagy receptor p62/SQSTM1 as an N-recognin whose activity is enhanced by Arg/N-degrons. Here, we show that Arg/N-degrons generated by *ATE1*-encoded R-transferase regulate p62-mediated mitophagy by promoting its recruitment to damaged mitochondria. Structural modification of Arg/N-degrons yielded ATB1071, a 443.5-Da orally bioavailable compound that activates p62 and induces stress-selective mitophagy through both Parkin-independent pathways involving NIPSNAP1 and NIPSNAP2, and a Parkin-dependent pathway involving the substrate EBP1/PA2G4. In *Ndufs4*^-/-^ mice, a Leigh syndrome (LS) model, ATB1071 induced mitophagy in the brain and exerted therapeutic benefits by reducing neuroinflammation, improving muscle strength and neuromuscular coordination, and extending lifespan. In cerebral ischemia-reperfusion (IR) model mice, ATB1071 reduced infarct volume and neuronal death, and ameliorated multiple behavioral deficits through EBP1-dependent mitophagy. Pharmacokinetic (PK) and toxicological analyses support ATB1071 as a preclinical candidate for mitochondria-associated neurological injury.

## INTRODUCTION

The mitochondrion serves as a central hub for cellular metabolism and signaling, playing essential roles in ATP production, metabolic homeostasis, and the regulation of cell death^1, 2^. Mitochondrial dysfunction, arising from genetic mutations or cellular stress is implicated in a wide range of diseases, including neurodegenerative disorders, diabetes, obesity, cardiomyopathies, aging-related conditions, and mitochondrial diseases^2–4^. Because damaged mitochondria can amplify oxidative stress, inflammation, and cell death, their selective removal is a central requirement for cellular homeostasis. To maintain mitochondrial integrity, cells rely on quality control systems that eliminate damaged mitochondria through the autophagy-lysosome system, a process referred to as mitophagy^2, 4^. Thus, the key therapeutic challenge is not simply to induce autophagy, but to activate mitochondrial clearance selectively at damaged organelles.

Mitophagy is initiated when normally short-lived PINK1 (PTEN-induced kinase 1) accumulates on depolarized mitochondria and recruits the E3 ligase Parkin^5, 6^. Parkin subsequently assembles Lys63-linked polyubiquitin chains on specific outer mitochondrial membrane (OMM) proteins, as well as on EBP1 (ErbB3-binding protein 1) which is translocated from the cytosol to the OMM upon mitochondrial damage^6–8^. These ubiquitin chains are recognized by the ubiquitin-binding domains (UBDs) of autophagy receptors, which in turn recruit LC3 anchored on autophagic membranes through their LC3-interacting region (LIR) domains, leading to lysosomal co-degradation^9–11^. Known mitophagy receptors include NDP52/CALCOCO2, NBR1 (neighbor of BRCA1 gene 1), p62/SQSTM1, TAX1BP1 (Tax1-binding protein 1), and OPTN/optineurin, which exhibit partially compensatory functions^10–12^. Among these, OPTN, NDP52, and, to a lesser extent, TAX1BP1 play dominant roles in mitophagy^13^, whereas p62 has often been viewed as a supporting receptor rather than a primary driver of mitochondrial clearance. However, we previously identified p62 as an autophagic N-recognin whose cargo-binding and self-assembly activities are allosterically activated by Arg/N-degrons, raising the possibility that p62 could be pharmacologically engaged to promote selective mitophagy^14, 15^.

Although Parkin-mediated ubiquitination is a major driver of mitophagy, several ubiquitin-independent mechanisms also recruit the autophagic machinery to damaged mitochondria.

Autophagic membranes containing LC3 can be directly tethered to impaired mitochondria through LIR domains present in OMM proteins such as BNIP3L/NIX^16^ and FUNDC1^17^. The inner mitochondrial membrane (IMM) protein Prohibitin 2 can likewise become exposed following proteasome-dependent rupture of the OMM, enabling conditional recruitment of LC3-positive phagophores^18^. More recently, the mitochondrial matrix proteins NIPSNAP1 and NIPSNAP2 were shown to localize to the surface of damaged mitochondria, where they recruit p62 or NDP52 independently of Parkin-mediated ubiquitination^19^. These studies indicate that damaged mitochondria can expose multiple receptor-engaging signals, but how p62 is selectively activated and assembled at these sites remains incompletely understood.

Mitochondrial diseases are a group of heterogeneous disorders caused by abnormal oxidative phosphorylation (OXPHOS), encompassing both primary mitochondrial diseases (PMD) and secondary mitochondrial dysfunction (SMD)^3, 4^. PMDs arise from pathogenic variants in either mtDNA or nuclear DNA that encode OXPHOS subunits, or in genes required for the assembly, maintenance, and regulation of the OXPHOS system^3, 20^. Among these disorders, Leigh syndrome (LS) is the most common pediatric PMD, with an estimated incidence of at least 1 in 40,000 newborns. LS is typically caused by mutations in *MT-ATP6*, *SURF1*, or *NDUFS4*^21, 22^. This severe neurometabolic condition is characterized by bilateral symmetrical lesions in the basal ganglia and/or brainstem, leading to respiratory failure and rapid deterioration of cognitive and motor functions^22, 23^. Beyond genetic mutations, SMD can also arise from pathological stressors that compromise mitochondrial functions, such as protein aggregates^24^, infectious pathogens^25^, and environmental toxins^26^. These insults can diminish ATP production, elevate reactive oxygen species (ROS), and trigger inflammatory responses driven by mtDNA leakage into the cytosol^27^. Thus, both inherited OXPHOS defects and acquired mitochondrial insults converge on damaged mitochondria as a pathogenic substrate that may be therapeutically removed by mitophagy.

Stroke is one of the leading causes of death worldwide^28^. Ischemic stroke accounts for approximately 85% of all stroke cases and arises from insufficient cerebral blood flow due to thrombotic or embolic vessel occlusion^28, 29^. Although restoring oxygen and vital nutrients is necessary to rescue threatened brain tissue, reperfusion paradoxically induces ischemia-reperfusion (IR) injury, which can amplify the initial ischemic damage^30^. Mitochondrial dysfunction is a central driver of reperfusion-associated stress and contributes to multiple pathological processes, including cell death, oxidative stress, inflammation, and breakdown of the blood-brain barrier (BBB)^30, 31^. While mitochondrial OXPHOS transiently recovers upon reperfusion, excessive Ca^2+^ influx and ROS promote opening of the mitochondrial permeability transition pore (mPTP), leading to ATP depletion and cell death^32^. Because mitochondrial impairment is an instigator of IR injury, mitochondria have become a major therapeutic target^31^. However, no approved pharmacological treatment directly removes damaged mitochondria or targets the mitochondrial dysfunction and neuronal death that drive IR-induced tissue injury^33^.

Over the past decades, natural products such as rapamycin^34^ and urolithin A^35^ have been identified as mitophagy inducers that modulate AMPK-mTOR signaling. Although these small molecules have demonstrated the therapeutic potential of autophagy induction in mitochondria-associated diseases, global mTORC1 inhibition may not be broadly advantageous, given mTOR’s pleiotropic role in regulating protein synthesis, immune function, and cell cycle progression^36–38^. To bypass mTOR-dependent mechanisms, small molecules were developed to directly recruit LC3^+^ phagophores to mitochondria, as exemplified by the synthetic BH3-mimetic compound UMI-77^39^ and the ATTEC (autophagy-tethering compounds) molecule mT1^40^, which enableMCL-1 (myeloid cell leukemia-1) to bind LC3 and tether LC3B to the OMM protein TSPO, respectively. While these approaches highlight the feasibility of directly linking mitochondria to the autophagy machinery, alternative strategies have focused on modulating the PINK1/Parkin pathway to enhance damage-specific signaling^41, 42^. Multiple PINK1/Parkin activators have been shown to induce mitophagy in the context of mitochondrial stress, yet candidates such as the PINK1 activator MTK458, whose derivative ABBV-1088 is currently in phase 1 clinical trials, inadvertently activated the mitochondrial integrated stress response (mitoISR) through a PINK1/Parkin-independent off-target mechanism^43^. Thus, current pharmacological strategies either broadly stimulate autophagy, artificially tether mitochondria to autophagic membranes, or modulate upstream damage-sensing pathways. A remaining challenge is to activate an endogenous autophagy receptor at damaged mitochondria without inducing mitochondrial stress or perturbing global cellular homeostasis.

The N-degron pathway safeguards the quality of both newly synthesized and mature proteins, as well as organelles and invading pathogens by directing them to proteasomal or lysosomal degradation^44–49^. In this system, single N-terminal amino acids of cytosolic and organellar proteins act as degradation signals known as N-degrons, which include type 1 residues (Arg, Lys, His; positively charged) and type 2 residues (Phe, Tyr, Trp, Leu, Ile; bulky hydrophobic)^44^. A key N-degron, N-terminal arginine (Nt-Arg), can be generated by *ATE1*-encoded R-transferases through conjugation of L-Arg to Nt-Glu, Nt-Asp, and oxidized Nt-Cys^44,50^. N-degrons are recognized by specific N-recognins, including the E3 ubiquitin ligases UBR1, UBR2, UBR4, UBR5, and KCMF1, which catalyze ubiquitination of their substrates^49, 51^. In addition to these ubiquitin ligases, we recently identified p62 as an autophagic N-recognin that binds Nt-Arg via its ZZ domain, triggering PB1 domain-mediated self-polymerization^14^.

Formation of filament-like p62 assemblies increases avidity for cargo through multivalent interactions, while simultaneously recruiting LC3^+^ phagophores through the exposed LIR domain^14, 15, 52, 53^. These findings raise the possibility that N-degron signals could directly activate p62 to promote selective autophagic clearance. To translate this N-degron principle into therapeutic strategies, we developed peptidomimetic small-molecule ligands that function as chemical N-degrons for the p62-ZZ domain, termed ATLs (autophagy-targeting ligands)^15^.

Previous studies from our group showed that chemical N-degrons can engage p62-dependent selective autophagy across diverse cargo classes, including protein aggregates, pathogens, lipid droplets, and damaged ER^15, 46, 54, 55^.

In this study, we identify the N-degron pathway as a key regulatory mechanism governing p62-mediated mitophagy. To therapeutically leverage this mechanism, we designed and synthesized an optimized chemical N-degron, ATB1071, which targets the p62-ZZ domain.

ATB1071 promoted the selective clearance of damaged mitochondria by engaging NIPSNAP1/2 and EBP1-dependent platforms for p62 recruitment, without activating AMPK-mTOR signaling. In *Ndufs4^-/-^* mice, a model of Leigh syndrome (LS), ATB1071 induced brain mitophagy, reduced neuroinflammation, improved neuromuscular function, and extended lifespan. In a cerebral ischemia-reperfusion (IR) injury model, ATB1071 conferred neuroprotection through EBP1-dependent mitophagy. Together, these findings establish chemical N-degrons as a therapeutic modality for the selective removal of damaged mitochondria via p62 modulation. The oral bioavailability, >65% brain penetration, and favorable preclinical safety profile of ATB1071 further support its development for mitochondria-associated diseases.

## RESULTS

### p62 mediates mitophagy as an N-recognin regulated by the N-degron pathway

Given our earlier findings that p62 functions as an N-recognin whose autophagic activity is allosterically regulated by N-degrons^14, 15^, we investigated whether this regulatory mechanism also operates during mitophagy (Fig. 1a). HeLa cells with minimal endogenous Parkin expression were engineered to stably express mt-mKeima^56^, a mitochondria-targeting fluorescent reporter whose excitation properties are pH-sensitive (Fig. 1b). Upon treatment with the mitochondrial uncoupler CCCP, which disrupts mitochondrial membrane potential (MMP), flow cytometry analyses revealed that *p62* knockdown led to an approximately 50% reduction in mitophagy, as measured by the ratio of 561-nm excitation in acidic lysosomes relative to 488-nm excitation at neutral pH (Supplementary Fig. 1a-d). A ∼38% reduction in mitophagy was also observed following *p62* knockdown in mt-mKeima HeLa cells ectopically expressing Parkin (Supplementary Fig. 1e, f). These results indicate that contributes substantially to mitophagy in both Parkin-independent and Parkin-dependent contexts.

**Figure 1.**
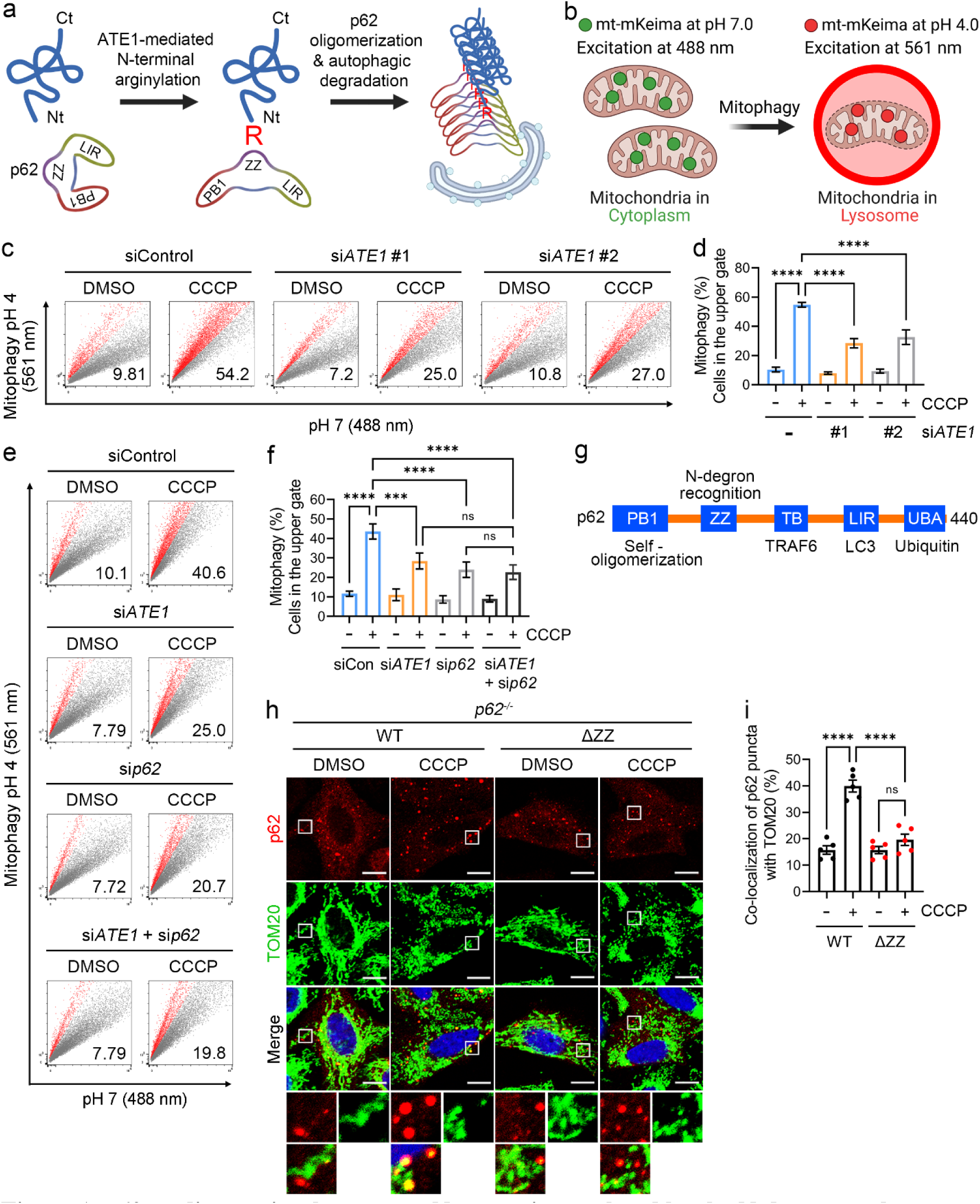
p62 mediates mitophagy as an N-recognin regulated by the N-degron pathway. **a** Depiction of the autophagic N-degron pathway. Ct, C-terminus; Nt, N-terminus; PB1, Phox and Bem1; ZZ, zinc finger; LC3-interacting region. **b** Schematic illustration of the mt-mKeima-based flow cytometry analysis used to quantify mitophagy. **c-f** Representative flow cytometry analyses of mitophagy in mt-mKeima HeLa cells. Cells were treated with CCCP (10 μM, 24 h) in the presence of the indicated siRNAs (**c**) and (**e**). Red dots and values indicate the population in the upper gate. Quantification of the upper-gated population is shown in (**d**) and (**f**) (*n* = 3). **g** Schematic diagram of the domain structure of p62. TB, TRAF6-binding; UBA, ubiquitin-associated. **h** Immunostaining of *p62*^-/-^ HeLa cells ectopically expressing wild-type (WT) p62 or p62 mutants lacking the ZZ domain. Cells were treated with CCCP (10 μM, 6 h). **i** Quantification of co-localization of p62 puncta with TOM20 is shown in (**h**). Insets (white boxes) are magnified below each panel. Scale bars, 10 μm (*n* = 5). Data are presented as means ± SD [(**d**) and (**f**)] or ± SEM (**i**). Statistical significances were calculated using one-way ANOVA. \*\*\**p* < 0.001; \*\*\*\**p* < 0.0001; ns, not significant.

We previously demonstrated that Arg/N-degrons bind the ZZ domain of p62 to promote its PB1-driven self-polymerization, a critical step for recruiting phagophores to sites of lysosomal degradation^14, 15^. To determine whether p62 mediates mitophagy through this N-recognin activity, we examined the role of ATE1, the enzyme responsible for generating Arg/N-degrons^14^.

Knockdown of *ATE1* reduced CCCP-induced mitophagy by ∼45% in mt-mKeima HeLa cells (Fig. 1c, d, Supplementary Fig. 1g, h), and co-depletion of ATE1 and p62 did not produce an additive effect (Fig. 1e, f). Thus, ATE1 and p62 appear to function in the same pathway during mitophagy. Consistent with an N-recognin–dependent mechanism, p62 mutants lacking either the ZZ or PB1 domain failed to translocate to mitochondria when ectopically expressed in CCCP-treated *p62*^-/-^ HeLa cells, as shown by co-immunostaining analyses of p62 and TOM20 (Fig. 1g-i, Supplementary Fig. 1i, j). These results indicate that p62 mediates mitophagy as an N-recognin whose function is activated through Arg/N-degron-dependent ZZ recognition and PB1-mediated self-assembly.

### ATB1071 activates p62 to induce mitophagy independently of AMPK–mTOR signaling

To develop mitophagy inducers that exploit the functional interplay between N-degrons and p62, we screened a library of ATL derivatives for their ability to activate p62 as an autophagic N-recognin. This initial screening identified YTK2205^54^ as a prototype compound capable of inducing mitophagy in mt-mKeima HeLa cells treated with CCCP (Fig. 2a, Supplementary Fig 2a, b). Guided by structure-activity relationship (SAR) analyses of its derivatives, we designed an orally available compound, ATB1071 (443.5 Da), featuring para-fluoro substituents on both benzyl ether groups and a propoxy tether connecting the catechol core to a secondary aminoethanol moiety, to improve potency and physicochemical properties (Fig. 2b, Supplementary Fig. 2c-e).

**Figure 2.**
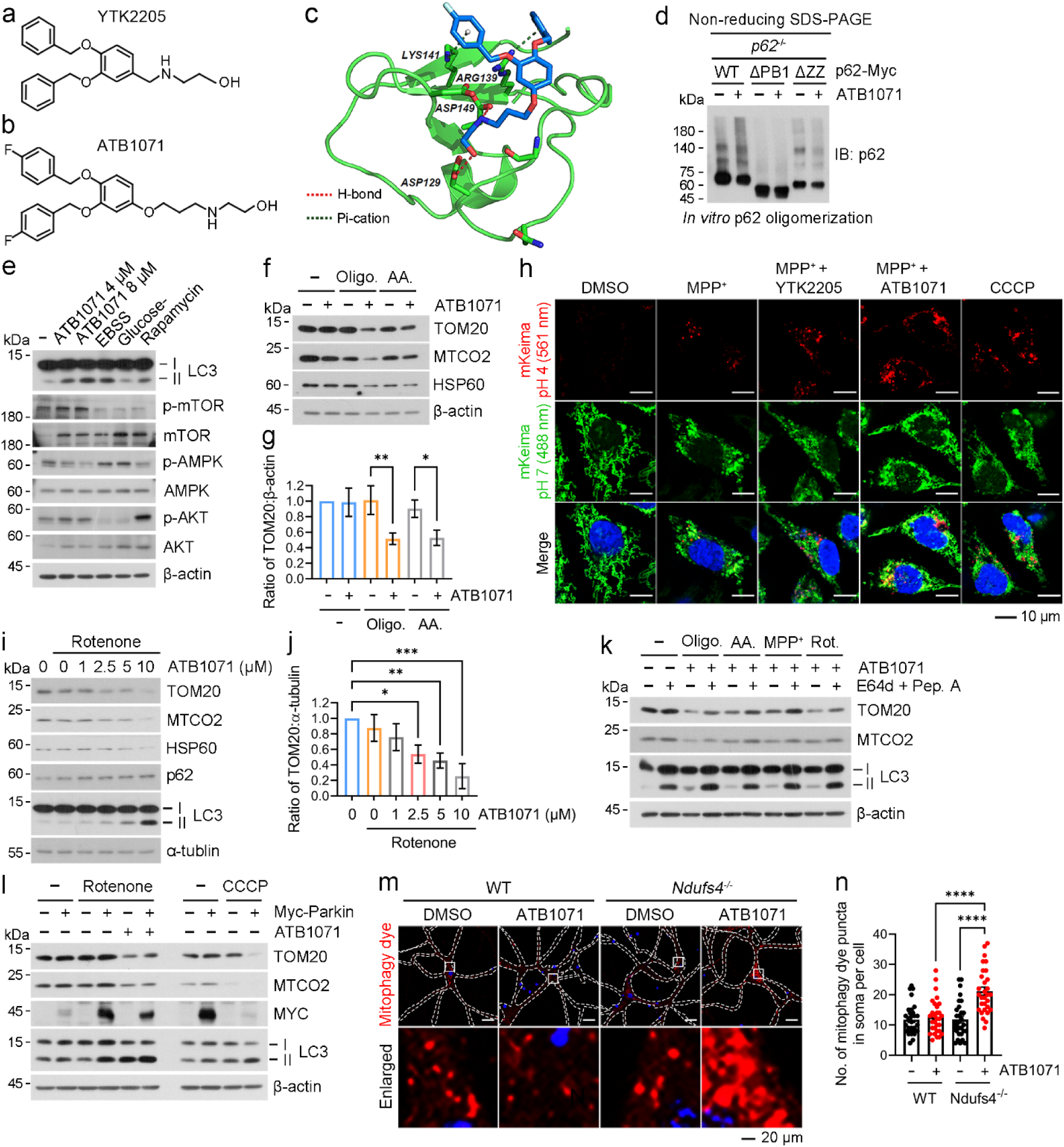
The chemical N-degron ATB1071 selectively induces mitophagy in damaged mitochondria independently of AMPK–mTOR signaling. **a**, **b** Structures of YTK2205 (**a**), and ATB1071 (**b**). **c** Three-dimensional docking model of ATB1071 bound to the ZZ domain of p62 showing key interacting residues and interactions. **d** *p62*^-/-^ HeLa cells ectopically expressing either WT p62 or the indicated p62 mutants were subjected to *in vitro* p62 oligomerization assay. **e** Immunoblotting analysis of SH-SY5Y cells the following treatment: ATB1071 (6 h), EBSS (6 h), glucose starvation (6 h), rapamycin (250 nM, 6 h). **f** Immunoblotting analysis of SH-SY5Y cells co-treated with ATB1071 (5 μM, 24 h) and the following mitochondrial stressors: oligomycin A (10 μM, 24 h), antimycin A (1 μM, 24 h). **g** Quantification of (**f**) (*n* = 3). **h** Representative images of mt-mKeima HeLa cells treated with CCCP (10 μM, 24 h), MPP^+^ (100 μM, 24 h), or the indicated combination treatments: MPP^+^ + YTK2205 (5 μM, 24 h) or MPP^+^ + ATB1071 (5 μM, 24 h). Scale bars, 10 μm. **i** Immunoblotting analysis of SH-SY5Y cells treated with the indicated concentrations of ATB1071 in the presence of rotenone (200 nM, 24 h). **j** Quantification of (**i**) (*n* = 3). **k** Immunoblotting analysis of SH-SY5Y cells co-treated with ATB1071 and the OXPHOS inhibitors in the presence of E64d + pepstatin A (10 mg/ml, 24 h). **l** Immunoblotting analyses of HeLa cells ectopically expressing Parkin following the indicated treatment: CCCP, rotenone, ATB1071, or a combination of rotenone and ATB1071. **m** Representative images of mitophagy dye staining in mouse primary neurons isolated from WT or *Ndufs4*^-/-^ mice treated with ATB1071 (4 μM, 24 h). Scale bars, 20 μm. **n** Quantification of (**m**) (*n* = 30). For (**n**), *n* values represent the number of cells from three biological replicates. Data are presented as means ± SD [(**g**) and (**j**)] or ± SEM (**n**). Statistical significances were calculated using one-way ANOVA. \**p* < 0.05; \*\**p* < 0.01; \*\*\**p* < 0.001; \*\*\*\**p* < 0.0001.

To assess whether ATB1071 engages the p62-ZZ domain, we modeled ATB1071 docking into the human p62-ZZ domain (PDB ID: 6MIU) using MAESTRO. The resulting 2D interaction diagram revealed that ATB1071 formed hydrogen bonds with Asp129 and Asp149 (within 3.5 Å) and π–cation interactions with Arg139 and Lys141 of the ZZ domain (Fig. 2c, Supplementary Fig. 2f). These residues overlap with the binding interface for the Arg-Glu peptide, supporting the idea that ATB1071 mimics an Arg/N-degron at the p62-ZZ domain^57^. We next evaluated the functional engagement of ATB1071 with the p62 ZZ domain. Under non-reducing SDS-PAGE conditions, ATB1071 induced p62 self-polymerization in a manner dependent on both the ZZ and PB1 domains (Fig. 2d). In *p62*^-/-^ HeLa cells reconstituted with wild-type p62 or mutant p62, ATB1071 promoted the formation of robust p62^+^ cytosolic puncta only in cells expressing wild-type p62, but not ZZ- or PB1-deficient mutants (Supplementary Fig. 2g, h). These puncta showed strong colocalization with LC3^+^ autophagic membranes (Supplementary Fig. 2i), indicating recruitment to autophagosomes. In SH-SY5Y neuroblastoma cells, ATB1071 enhanced LC3 lipidation (LC3-I to LC3-II conversion) without increasing phosphorylation of AMPKα at Thr172 or decreasing phosphorylation of mTOR at Ser2448 (Fig. 2e). Thus, ATB1071 activates p62-dependent selective autophagy without engaging canonical AMPK activation or mTOR inhibition. Collectively, these findings demonstrate that ATB1071 functions as a chemical N-degron that engages the p62-ZZ domain, promotes PB1-dependent p62 assembly, and activates selective autophagy through a noncanonical mechanism.

### ATB1071 selectively induces mitophagy in damaged mitochondria

Because several previously reported mitophagy inducers act even under basal, non-stress conditions^35, 39^, we first evaluated ATB1071 in SH-SY5Y cells maintained under normal culture conditions. We monitored the turnover of TOM20, MTCO2, and HSP60, which mark the mitochondrial outer membrane, inner membrane, and matrix, respectively. ATB1071 treatment did not produce appreciable changes in the abundance of these proteins (Supplementary Fig. 3a). Thus, ATB1071 has minimal mitophagy activity in the absence of mitochondrial stress. In contrast, ATB1071 robustly promoted the degradation of these mitochondrial markers when mitochondria were damaged with inhibitors of the OXPHOS complexes, including oligomycin A, antimycin A, rotenone, and MPP^+^ (1-methyl-4-phenylpyridinium) (Fig. 2f, g, Supplementary Fig. 3b, c). Flow cytometry confirmed that this accelerated mitochondrial turnover corresponded with increased lysosomal targeting of mt-mKeima (Supplementary Fig. 3d, e). Similar mitophagy responses were observed with YTK2205, the precursor scaffold of ATB1071, as shown by live-cell confocal imaging (Fig. 2h, Supplementary Fig. 3f) and flow cytometry (Supplementary Fig. 3g, h). ATB1071 exhibited dose-dependent activity at 2.5-10 μM in immunoblotting analyses (Fig. 2i, j). WST viability assays yielded IC_50_ values of 14.75 μM in SH-SY5Y cells and 11.51 μM in HeLa cells (Supplementary Fig. 3i, j). Importantly, ATB1071 failed to promote mitochondrial marker degradation when p62 was depleted or when lysosomal proteases were inhibited with E64d plus pepstatin A (Fig. 2k, Supplementary Fig. 3k). These results show that ATB1071 induces p62-dependent, autophagy-lysosome-mediated mitochondrial clearance selectively under mitochondrial stress.

Next, we examined whether ATB1071 induces mitophagy through the autophagic N-degron pathway. In HeLa cells treated with rotenone, ATB1071 accelerated the colocalization of p62 with TOM20 in immunostaining analyses (Supplementary Fig. 4a, b) and promoted the mitochondrial translocation of p62 in fractionation analyses (Supplementary Fig. 4c, d). This p62 recruitment preceded or accompanied the assembly of LC3^+^ phagophores on TOM20^+^ mitochondria, an event that was absent in *p62^-/-^* HeLa cells (Supplementary Fig. 4e, f). Thus, ATB1071-induced mitophagy requires p62 to recruit autophagic membranes to damaged mitochondria. We next assessed the role of Parkin in this process. Immunoblotting showed that ATB1071 did not exhibit synergistic degradation activity when combined with Parkin overexpression under 200 nM rotenone treatment—an experimental condition that does not elicit Parkin-mediated mitophagy, in contrast to CCCP (Fig. 2l) indicating that ATB1071 can act independently of Parkin under this stress condition. By contrast, in CCCP-treated cells, ATB1071 further augmented Parkin-mediated mitophagy, as measured by flow cytometry (Supplementary Fig. 4g, h). Together, these findings indicate that chemical N-degrons allosterically activate p62, enabling its self-polymerization on the outer membrane of damaged mitochondria and the subsequent recruitment of LC3^+^ phagophores for lysosomal co-degradation.

### ATB1071 alleviates mitochondrial dysfunction under genetic and chemical mitochondrial stress

Mutations in *NDUFS4* produce the most severe clinical phenotype of LS among OXPHOS complex I-related defects^58^. We therefore evaluated whether ATB1071 can restore mitochondrial quality control in LS-relevant cellular models. In *NDUFS4* knockdown SH-SY5Y cells, ATB1071 induced oligomeric and aggregated species of p62 in the mitochondrial fraction (Supplementary Fig. 5a-d). The mitochondrial recruitment of p62 was associated with increased lysosomal delivery of mitochondria, as observed by live-cell confocal imaging of mt-mKeima HeLa cells (Supplementary Fig. 5e). Similar mitophagy activation was obtained in primary neurons derived from *Ndufs4*^-/-^ murine brains (Fig. 2m, n). Dose-dependent activity was detected at 2.5-10 μM in immunoblotting analyses of primary neurons (Supplementary Fig. 5f, g) and at 2-8 μM by flow cytometry of mt-mKeima Hela cells (Supplementary Fig. 5h). These results indicate that ATB1071 activates mitophagy in cellular models of OXPHOS complex I deficiency.

Dysregulation of OXPHOS complex I lowers MMP and elevates ROS, thereby diminishing the proton-motive force required for ATP synthase and ultimately impairing ATP production^59–61^. We therefore assessed whether ATB1071 functionally rescues these mitochondrial defects. In *NDUFS4* knockdown cells, ATB1071 restored the depolarized MMP to near normal levels (Fig. 3a, b). Comparable restoration of MMP was observed in cells undergoing chemically induced mitochondrial damage via CCCP treatment, an effect that was attenuated by the autophagy inhibitor bafilomycin A1 (Fig. 3c, d), linking functional rescue to autophagy-dependent mitochondrial turnover. Consistently, ATB1071 normalized the elevated intracellular ROS levels induced by either *NDUFS4* knockdown or rotenone exposure (Fig. 3e-h). Moreover, the ∼20% reduction in ATP levels caused by *NDUFS4* knockdown was fully rescued by ATB1071 treatment (Fig. 3i). Extracellular flux analyses further revealed that ATB1071 increased mitochondrial ATP production and basal respiration—both diminished by *NDUFS4* knockdown—while maximal respiration was slightly increased but did not reach statistical significance (Fig. 3j-m). Thus, ATB1071 not only activates mitophagy but also restores mitochondrial bioenergetic function under genetic and chemical mitochondrial stress.

**Figure 3.**
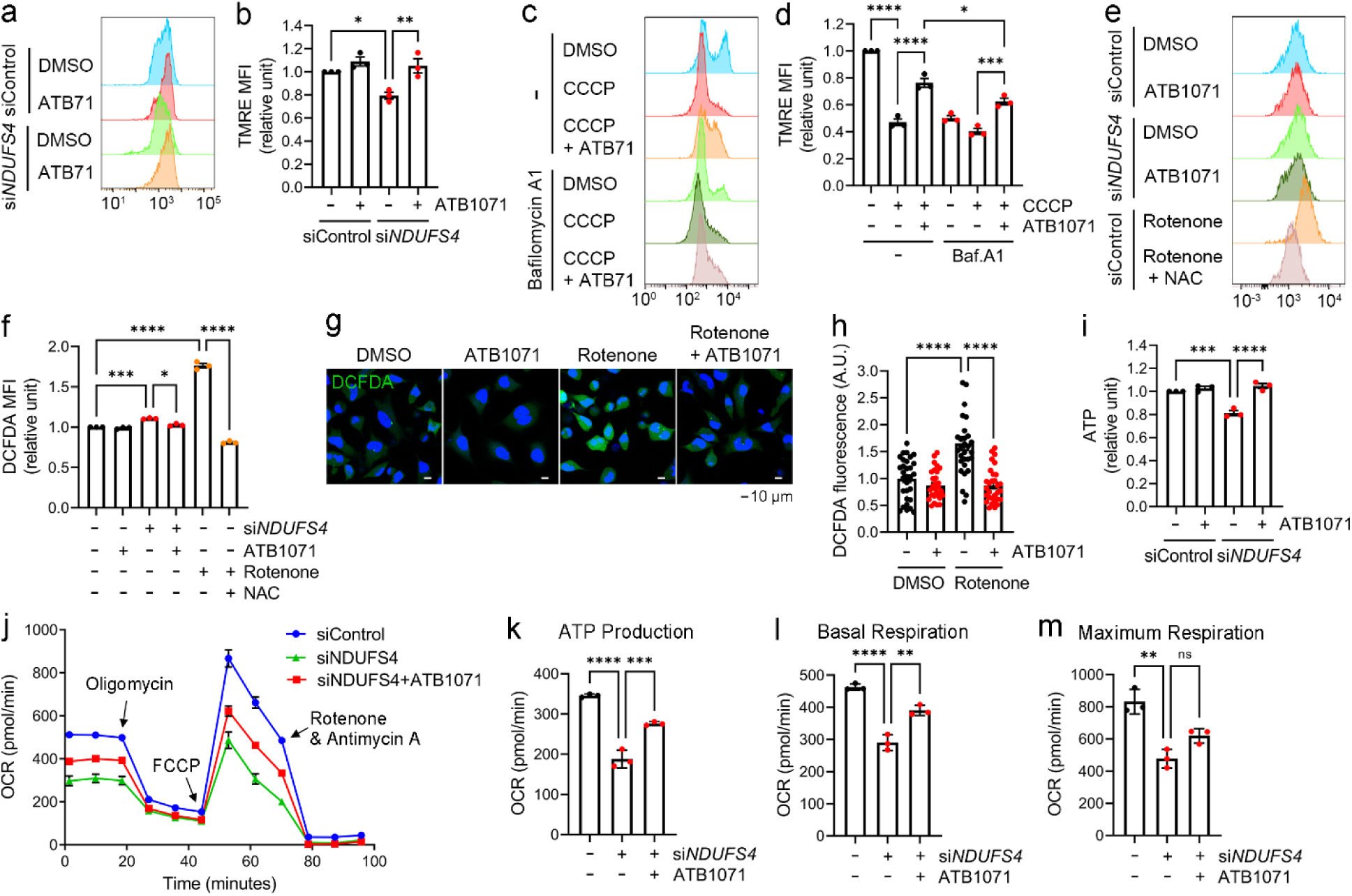
ATB1071 alleviates mitochondrial dysfunction under genetic and chemical mitochondrial stress. **a-f** Flow cytometry of SH-SY5Y cells with the indicated treatments: ATB1071 (5 μM, 24 h), rotenone (200 nM, 24 h), NAC (1 mM, 24 h), CCCP (20 μM, 6 h), and bafilomycin A1 (200 nM, 24h). Quantification of geometric mean fluorescence intensity (MFI) of TMRE (**b**) and (**d**) or DCFDA (**f**) (*n* = 3). **g** DCFDA staining of HeLa cells co-treated with rotenone (200 nM, 24 h) and ATB1071 (5 μM, 24 h). **h** Quantification of (**g**) (*n* = 30). **i** ATP levels of SH-SY5Y cells treated with ATB1071 (5 μM, 24 h) in the presence of si*NDUFS4* (*n* = 3). **j-m** Oxygen consumption rate (OCR) of SH-SY5Y cells treated with ATB1071 in the presence of si*NDUFS4.* Quantification of basal respiration (**k**), mitochondrial ATP production (**l**), and maximal respiration (**m**) is shown (*n* = 3). For (**h**), *n* values represent the number of cells from three biological replicates. Data are presented as means ± SEM. Statistical significances were calculated using one-way ANOVA. \**p* < 0.05; \*\**p* < 0.01; \*\*\**p* < 0.001; \*\*\*\**p* < 0.0001.

### NIPSNAP1 and NIPSNAP2 recruit p62 to damaged mitochondria during ATB1071-induced mitophagy

To identify mitochondrial adaptors that recruit ATB1071-activated p62 during mitophagy, we performed mass spectrometric analyses of p62 immunoprecipitates from ATB1071-treated HeLa cells. Of the 122 mitochondrial proteins identified in the p62 interactome, 31 were increased upon ATB1071 treatment (Fig. 4a, b). Among these candidates, NIPSNAP1 and NIPSNAP2 have been reported to act as “eat-me” signals that accumulate on the OMM to recruit autophagy receptors, including p62 and NDP52^19^. Co-immunoprecipitation (Co-IP) validated that ATB1071 induced p62 binding to NIPSNAP1 in a dose-dependent manner (Fig. 4c, d), and this interaction was further enhanced when mitochondria were damaged by CCCP (Fig. 4e, f). Flow cytometry of *NDUFS4* knockdown mt-mKeima HeLa cells demonstrated that ATB1071-induced mitophagy was suppressed upon knockdown of either *NIPSNAP1* or *NIPSNAP2*, but not *CLS* (cardiolipin synthase), *PLSCR3* (phospholipid scramblase 3)^62^, or *RBX1* (ring-box 1)^63^ (Supplementary Fig. 6a, b). Consistently, simultaneous knockdown of *NIPSNAP1* and *NIPSNAP2* led to an ∼53% reduction in mitophagy induced by ATB1071 and CCCP co-treatment (Fig. 4g, h). Thus, NIPSNAP1 and NIPSNAP2 are functionally required for efficient ATB1071-induced mitophagy. We next examined whether NIPSNAP1 and NIPSNAP2 are exposed to the surface of damaged mitochondria. In CCCP-treated SH-SY5Y cells and *Ndufs4*^-/-^ mouse brains, NIPSNAP1 and NIPSNAP2 remained in the pellet after alkaline sodium carbonate (Na_2_CO_3_) extraction, and were readily digested by proteinase K, indicating their localization to the OMM surface (Supplementary Fig. 6c-f). These results show that NIPSNAP1 and NIPSNAP2 form damage-exposed OMM platforms that recruit p62 during ATB1071-induced mitophagy.

**Figure 4.**
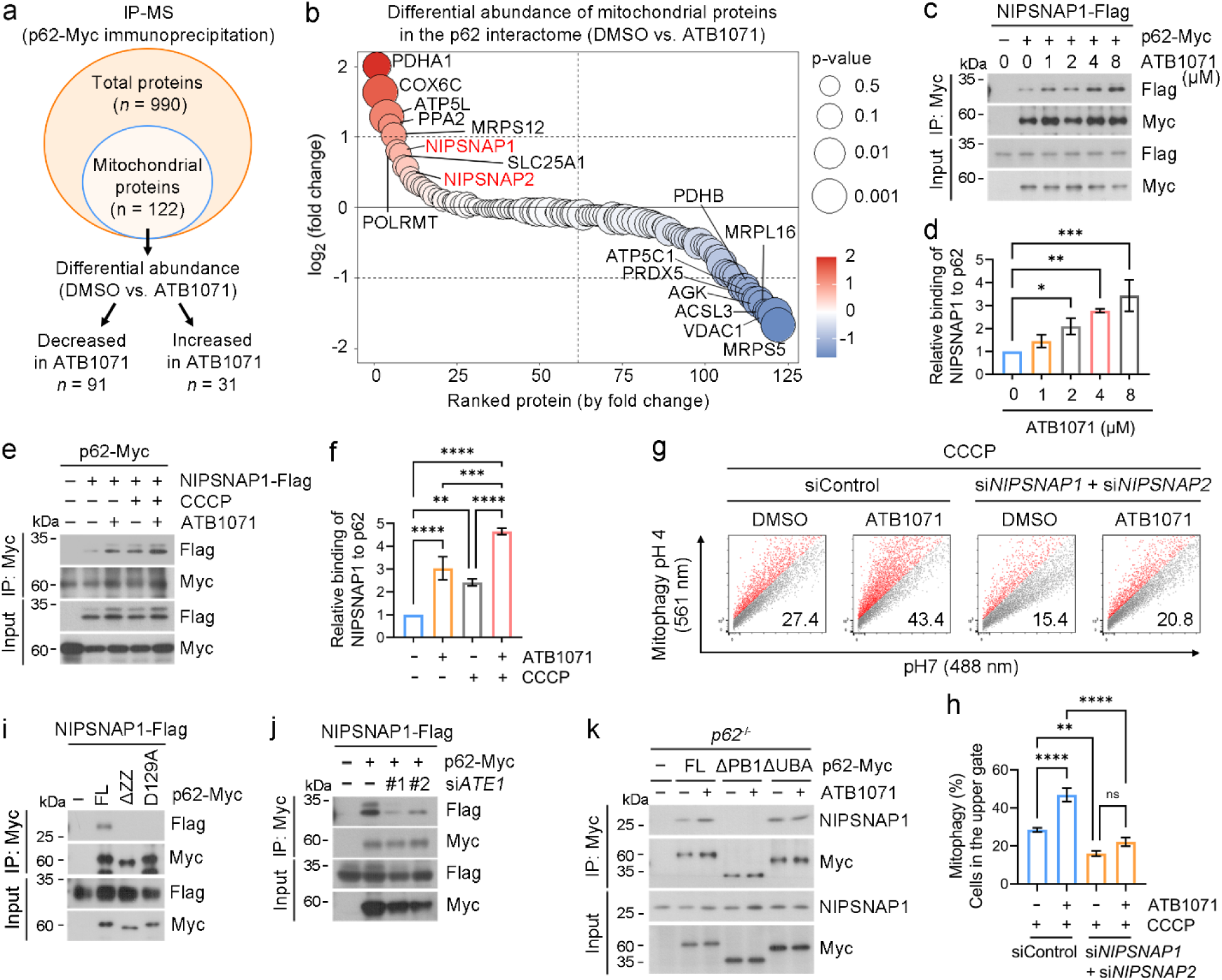
NIPSNAP1 and NIPSNAP2 recruit p62 to damaged mitochondria during ATB1071-induced mitophagy. **a** Classification of proteins identified by IP-MS in vehicle- or ATB1071-treated HeLa cells (2 μM, 24 h) (*n* = 3). **b** Rank-ordered plot of differential abundance of mitochondrial proteins upon ATB1071 treatment. Proteins are ranked by log_2_ fold change with color and point size indicating log_2_ fold change and p-value, respectively. **c** Co-IP analysis of 293T cells ectopically expressing NIPSNAP1-Flag and p62-Myc following ATB1071 treatment at the indicated concentration for 8 h. **d** Quantification of (**c**) (*n* = 3). **e** Co-IP analysis of 293T cells expressing NIPSNAP1-Flag and p62-Myc. Cells were treated with CCCP (10 μM, 8 h), ATB1071 (5 μM, 8 h), or a combination of CCCP and ATB1071. **f** Quantification of (**e**) (*n* = 3). **g** Representative flow cytometry analyses of mitophagy in mt-mKeima HeLa cells. Cells were treated with either CCCP (10 μM, 24 h), ATB1071, or a combination of CCCP and ATB1071 in the presence of si*NIPSNAP1* and si*NIPSNAP2*. **h** Quantification of (**g**) (*n* = 3). **i**, **j** Co-IP analysis assessing the binding between NIPSNAP1-Flag and various p62-Myc mutants (**i**), the effects of ATE1 siRNAs on NIPSNAP1-Flag/p62-Myc interaction (**j**). **k** Co-IP analysis of *p62*^-/-^ HeLa cells ectopically expressing either WT p62 or the indicated p62 mutants. Cells were treated with ATB1071 (5 μM, 8 h). Data are presented as means ± SD. Statistical significances were calculated using one-way ANOVA. \**p* < 0.05; \*\**p* < 0.01; \*\*\**p* < 0.001; \*\*\*\**p* < 0.0001; ns, not significant.

As an N-recognin, p62 translocates to mitochondria and mediates mitophagy (Fig. 1).

To determine whether p62 binds NIPSNAP1 and NIPSNAP2 through its N-recognin module, we mapped the domains required for binding (Supplementary Fig. 1j). Co-IP analyses showed that their interactions were abolished by the deletion of the ZZ domain (Fig. 4i). Consistent with the docking model, in which ATB1071 formed hydrogen bonding with Asp129 (Fig. 2c, Supplementary Fig 2f), the D129A mutation was sufficient to disrupt p62 binding to NIPSNAP1/2 (Fig. 4i, Supplementary Fig. 6g). These interactions were also impaired in *ATE1* knockdown HEK293T cells and *ATE1^-/-^* HeLa cells (Fig. 4j, Supplementary Fig. 6h). In addition to the ZZ domain, optimal binding required the PB1 domain that mediates p62 self-polymerization and cargo clustering (Fig. 4k). By contrast, deletion of the UBA domain did not impair p62-NIPSNAP1 binding (Fig. 4k), consistent with previous reports that Parkin does not ubiquitinate NIPSNAP1/2^19^. Thus, p62 engagement of NIPSNAP1/2 depends on the N-degron-sensitive ZZ/PB1 axis rather than the UBA-dependent ubiquitin-binding module. As a chemical Arg/N-degron, ATB1071 substantially restored p62-NIPSNAP1 interactions in ATE1-deficient cells but not in cells lacking the p62 ZZ or PB1 domains (Fig. 4k, Supplementary Fig. 6i, j). Moreover, ATB1071 promoted the formation of NIPSNAP1^+^ puncta that prominently colocalized with p62^+^ puncta (Supplementary Fig. 6k). Collectively, these results suggest that ATB1071 drives the assembly of p62 polymers that establish multivalent interactions with NIPSNAP1/2 exposed on the OMM of damaged mitochondria, thereby marking damaged mitochondria for autophagic clearance.

### ATB1071 mitigates neuroinflammation and pathological phenotypes in *Ndufs4*^-/-^ mice

To determine whether ATB1071 activates mitophagy in vivo, *Ndufs4^-/-^* mice, an LS model with OXPHOS complex I deficiency, were intraperitoneally injected with 10 mg/kg ATB1071 every other day (QAD) for 40 days beginning at postnatal day 10 (Fig. 5a).

**Figure 5.**
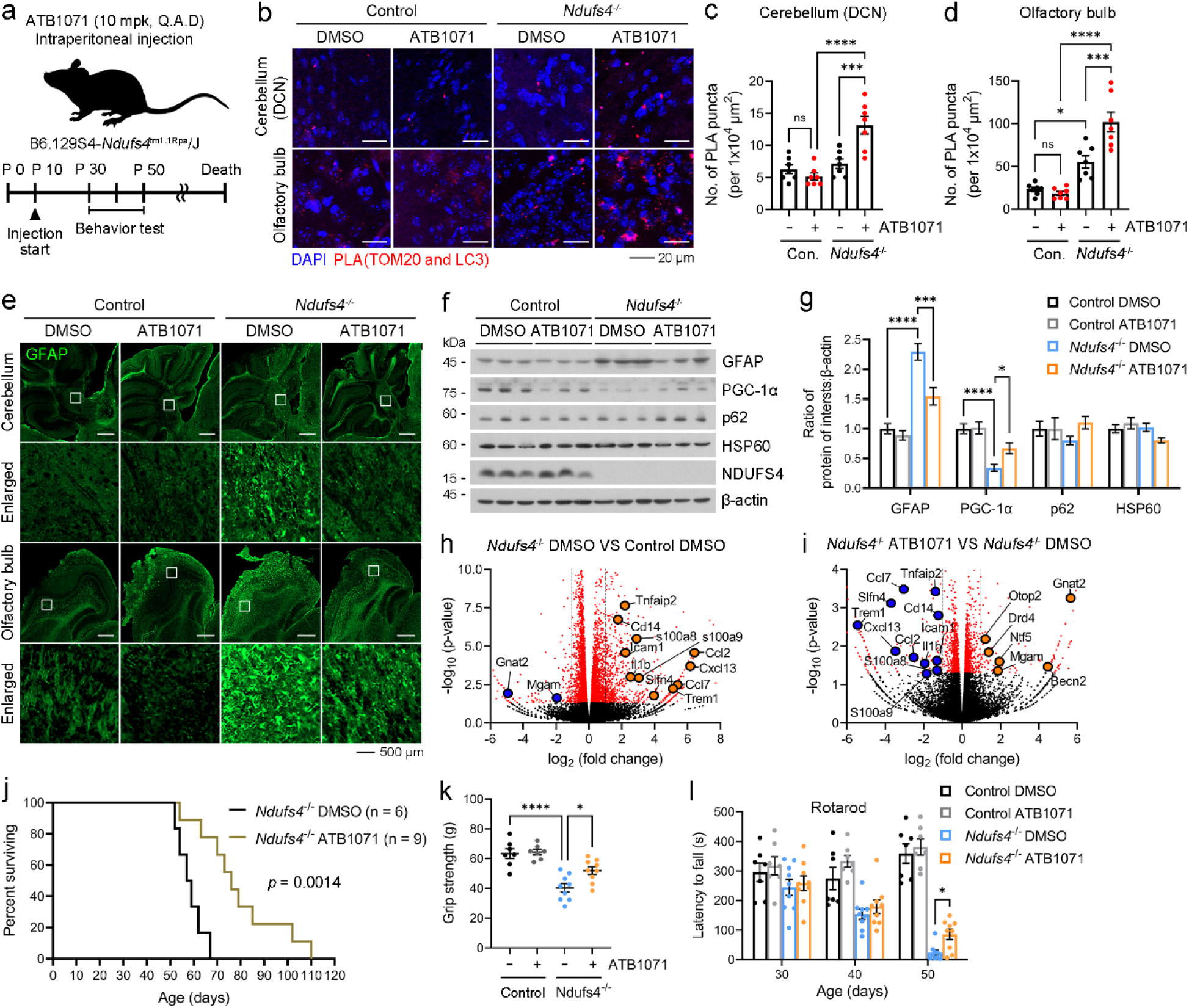
ATB1071 induces mitophagy and mitigates pathological phenotypes in *Ndufs4*^-/-^ mice. **a** Schematic illustration of the injection timeline and experimental timeline. **b-d** Proximity ligation assay (PLA) performed in the deep cerebellar nuclei (DCN) and olfactory bulb of control or *Ndufs4*^-/-^ mice (**b**). Quantification of PLA puncta in the DCN and olfactory bulb is shown in (**c**) and (**d**), respectively. Scale bars, 20 μm (*n* = 7 mice). **e** GFAP staining of the cerebellum and olfactory bulb from control and *Ndufs4*^-/-^ mice. Regions within white boxes are shown at higher magnification below. Scale bars, 500 μm (*n* = 8 mice). **f** Immunoblotting analyses of whole brains lysates from *Ndufs4*^-/-^ mice using the indicated antibodies. **g** Quantification of (**f**) (*n* = 12 mice). **h**, **i** Volcano plots of differentially expressed genes (DEGs) in the following comparisons: vehicle-treated control versus vehicle-treated *Ndufs4*^-/-^ mice (**h**), and vehicle-treated versus ATB1071-treated *Ndufs4*^-/-^ mice (**i**). Vertical dashed lines indicate fold-change thresholds; horizontal dashed lines indicate a *p*-value threshold of 0.05 (*n* = 3 mice). **j** Survival curves of *Ndufs4*^-/-^ mice treated with DMSO or ATB1071 (*n* = 6, *n* = 9, respectively). **k** Fore limbs grip strength measured for control at postnatal day 40 (P40) (*n* = 7-9 mice). **l** Accelerated rotarod performance measured at the indicated ages (*n* = 7-9 mice). For (**k**) and (**l**), median values of triplicate trials per mouse are shown. Data are presented as means ± SEM. Statistical significances were calculated using one-way ANOVA [(**c**), (**d**), and (**k**)], Log-rank test (**j**), or two-way ANOVA (**l**). \**p* < 0.05; \*\**p* < 0.01; \*\*\**p* < 0.001; \*\*\*\**p* < 0.0001; ns, not significant.

Brain sections were subjected to proximity ligation assays (PLA), which visualize *in situ* protein-protein interaction within ∼40 nm. ATB1071 treatment markedly increased TOM20^+^LC3^+^ PLA puncta in both the deep cerebellar nuclei (DCN) and olfactory bulb of *Ndufs4*^-/-^ mice relative to vehicle-treated controls (Fig. 5b-d). Fractionation of *Ndufs4*^-/-^brain extracts revealed that ATB1071 elevated p62 levels by ∼1.8-fold and its oligomeric/aggregated species by ∼2.4-fold in the mitochondrial fraction (Supplementary Fig. 7a-d). These results indicate that ATB1071 enhances p62-associated mitophagy in *Ndufs4*^-/-^ mouse brains.

The lethality of *Ndufs4*^-/-^ mice is closely associated with neuroinflammation, characterized by astrocyte activation and glial reactivity in the cerebellum, vestibular nuclei, and olfactory bulb^64, 65^. To determine whether ATB1071 attenuates neuroinflammation, we performed immunostaining for GFAP (glial fibrillary acidic protein) and Iba1 (ionized calcium-binding adapter molecule 1), markers of astrocytes and microglia, respectively. ATB1071 administration significantly reduced GFAP^+^astrocytes and Iba1^+^ microglia in *Ndufs4*^-/-^ brains (Fig. 5e, Supplementary Fig. 8a, b). Immunoblotting confirmed reduced GFAP protein levels following ATB1071 treatment (Fig. 5f, g). Transcriptome sequencing of wild-type and *Ndufs4*^-/-^ brains revealed 865 differentially expressed genes (DEGs), of which 703 were upregulated and 162 were downregulated (Supplementary Fig. 8c). Gene Ontology (GO) analyses showed enrichment of immune- and inflammation-related biological processes in *Ndufs4*^-/-^ brains, whereas no significant GO terms were identified among downregulated genes (Supplementary Fig. 8d). Notably, GO analyses of genes downregulated by ATB1071 in *Ndufs4*^-/-^ brains revealed enrichment of biological processes associated with immune cell recruitment and inflammation, while genes upregulated by ATB1071 were enriched for homeostatic processes (Supplementary Fig. 8e, f). ATB1071 also lowered the expression of specific inflammation-associated genes in *Ndufs4*^-/-^ brains, including Cd14, Il-lβ (interleukin-1 beta), and S100a8, as compared with vehicle-treated *Ndufs4*^-/-^ brains (Fig. 5h, i). Such effects were not observed in wild-type mice treated with ATB1071 (Supplementary Fig. 8g).

These results indicate that ATB1071 reverses the inflammatory transcriptional state and suppresses glial activation caused by OXPHOS complex I dysfunction in *Ndufs4*^-/-^ brains.

We next assessed the therapeutic impact of ATB1071 on disease manifestations in *Ndufs4*^-/-^ mice. Body weight trajectories did not differ significantly over the entire study period (Supplementary Fig. 9a). However, ATB1071 extended median survival by ∼31% and maximal lifespan by ∼64% relative to vehicle-treated *Ndufs4*^-/-^ mice (Fig. 5j). In the forelimb grip strength test, ATB1071-treated *Ndufs4*^-/-^ mice exhibited ∼34% improvement after 30 days of treatment, an effect not observed in control mice (Fig. 5k). ATB1071 also mitigated progressive neuromuscular decline, with treated *Ndufs4*^-/-^ mice exhibiting a ∼3-fold increase in rotarod latency to fall at postnatal day 50 compared with vehicle-treated counterparts (Fig. 5l). Additionally, the onset of clasping/twisting—a hallmark of neurological deterioration^64^—was delayed by ∼12% with ATB1071 (Supplementary Fig. 9b). To assess the impact of ATB1071 on skin inflammation-associated hair loss around weaning^65^, *Ndufs4*^-/-^ mice were scored at postnatal day 20 on a 0-2 severity scale (Supplementary Fig. 9c). ATB1071 significantly reduced the mean score from 2 to 1.22, indicating an anti-inflammatory effect during the early treatment window (postnatal days 10-20; Supplementary Fig. 9d). These results demonstrate that ATB1071 engages brain mitophagy, suppresses neuroinflammation, and improves survival and neuromuscular function in *Ndufs4*^-/-^ mice.

### ATB1071 provides neuroprotection and restores functional performance through Parkin-dependent mitophagy following IR injury

Our previous work demonstrated that, during mitochondrial damage caused by IR injury, EBP1 is translocated to the surface of dysfunctional mitochondria and ubiquitinated by Parkin, thereby recruiting p62 to initiate mitophagy^8, 66^. To determine whether ATB1071 confers therapeutic benefits in cerebral IR injury through EBP1-dependent mitophagy, we generated neuron-specific *Ebp1* knockout mice (*CamK*Ⅱ*-Cre*; *Pa2g4*^flox/flox^; hereafter referred to as *Ebp1* CKO). Cerebral IR injury was induced in wild-type and *Ebp1* CKO mice by transient middle cerebral artery occlusion (MCAO), followed by intraperitoneal administration of 10 mg/kg ATB1071 at 0, 3, 24, and 48 h post-MCAO (Fig. 6a). Immunoblotting analyses showed that ATB1071 decreased TOM20 levels while increasing LC3 and p62 in the hippocampus of MCAO-subjected control mice, indicating activation of mitophagy after IR injury (Fig. 6b, c). This effect was markedly diminished in MCAO-subjected *Ebp1* CKO mice (Fig. 6b, c). Consistently, immunostaining revealed that ATB1071-driven recruitment of LC3^+^ phagophores to TOM20^+^ mitochondria was substantially impaired in *Ebp1* CKO mice compared with controls (Fig. 6d). These data indicate that ATB1071 accelerates mitophagy during cerebral IR injury largely through the EBP1–p62 axis.

**Figure 6.**
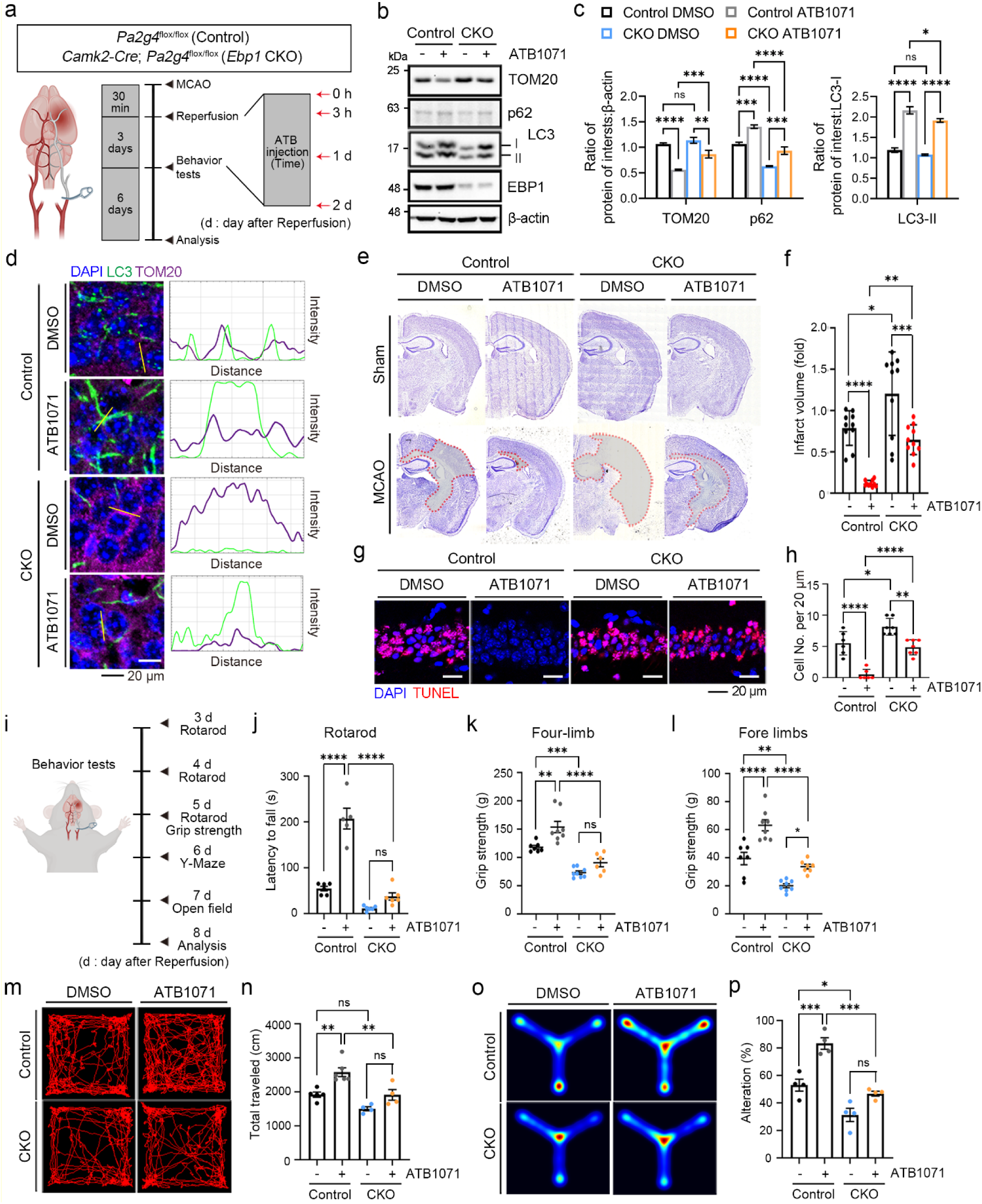
ATB1071 provides neuroprotection and restores functional performance through Parkin-dependent mitophagy following IR injury. **a** Schematic illustration of the experimental design. **b** Immunoblot analyses of hippocampal lysates of MCAO mice using the indicated antibodies. **c** Quantification of (**b**) (*n* = 5 mice). **d** Immunostaining of hippocampal tissue from MCAO mice. The fluorescence intensity profiles along the yellow lines are displayed on the right. Scale bar, 20 µm. **e** Nissl staining of brain sections from the indicated groups. **f** Quantification of (**e**) (*n* = 10 mice). **g** TUNEL staining of the hippocampal CA1 region. Scale bar, 20 µm. **h** Quantification of (g) (*n* = 6-7 mice). **i** Schematic illustration of the behavioral testing paradigm. **j-n** Motor-related behavioral assessments, including rotarod performance (**j**), four-limb grip strength (**k**) and forelimb grip strength (**l**), and open-field exploration (**m**) and (**n**) (*n* = 4-8 mice). **o**, **p** Y-maze spontaneous alternation test. The percentage of arm alteration was measured and calculated (*n* = 4 mice). For (**j**), (**k**) and (**l**), median values of triplicate trials per mouse are shown. Data are presented as mean ± SEM. Statistical significances were calculated using one-way ANOVA. \**p* < 0.05; \*\**p* < 0.01; \*\*\**p* < 0.001; \*\*\*\**p* < 0.0001; ns, not significant.

To assess the neuroprotective effects of ATB1071 and its dependence on EBP1, we determined whether ATB1071 attenuates brain damage caused by MCAO. As expected, *Ebp1* CKO mice exhibited significantly larger infarct volumes after IR compared with control mice, confirming that loss of *Ebp1* increases susceptibility to ischemic damage (Fig. 6e, f). Notably, ATB1071 administration reduced infarct volume in control mice by ∼85% (Fig. 6e, f). This protective efficacy, however, was markedly blunted in *Ebp1* CKO mice, in which ATB1071 reduced infarct volume by only ∼46%. (Fig. 6e, f). Consistently, TUNEL staining revealed that ATB1071 lowered neuronal death in the CA1 region of MCAO-subjected control mice by ∼91%, whereas the reduction observed in ATB1071-treated *Ebp1* CKO mice was limited to ∼40% (Fig. 6g, h). These results demonstrate that ATB1071 confers robust neuroprotection following cerebral IR injury, and that this efficacy relies predominantly on EBP1-mediated mitophagy.

We also evaluated the therapeutic efficacy of ATB1071 on neurological function and its dependence on EBP1 (Fig. 6i). In the rotarod test, ATB1071-treated control mice showed a robust improvement in motor coordination, with latency to fall increasing from 54.7 to 207.4 seconds. In contrast, ATB1071-treated *Ebp1* CKO mice displayed no significant improvement compared with vehicle-treated mutants (Fig. 6j). Similarly, ATB1071 enhanced four-limb and forelimb grip strengths in MCAO-subjected control mice by ∼1.3-fold and ∼1.6-fold, respectively, whereas *Ebp1* CKO mice exhibited minimal or no enhancement (Fig. 6k, l). Open-field behavioral analyses further revealed that ATB1071 improved ambulatory activity in MCAO-subjected control mice, increasing total travel distance (from 1919 to 2579 cm) and promoting exploration of the center area of the maze (Fig. 6m, n). By contrast, ATB1071-treated *Ebp1* CKO mice exhibited no significant improvement and continued to display wall-hugging behavior (Fig. 6m, n). In the Y-maze test, ATB1071 markedly rescued working memory deficits in control mice, raising the spontaneous alternation rate from 53% to 83% (Fig. 6o, p). However, these cognitive benefits were substantially reduced in *Ebp1* CKO mice, in which alternation increased only from 31% to 47% (Fig. 6o, p). These results demonstrate that ATB1071 significantly improves motor, behavioral, and cognitive functions after cerebral IR injury, and that these therapeutic benefits are largely dependent on EBP1-mediated mitophagy.

### EBP1 reintroduction restores ATB1071-mediated mitophagy and neuroprotection after ischemic injury

Given our finding that EBP1 is required for ATB1071-induced mitophagy and neuroprotection in MCAO mice, we evaluated whether reintroducing EBP1 could restore ATB1071 responsiveness in the ischemic brain. To reintroduce EBP1, AAV-*Ebp1* or the control AAV-mock was stereotaxically injected into the prospective ischemic region of *Ebp1* CKO mice five days before MCAO induction. Immunoblotting analyses showed that ATB1071 treatment restored mitophagy activation in EBP1-reintroduced *Ebp1* CKO mice, as evidenced by increased LC3 and p62 (Fig. 7a, b). The ATB1071-induced increase in LC3 was associated with increased mitochondrial localization of LC3 in the hippocampus, as determined by immunostaining (Fig. 7c). Consistent with restored mitophagy activation, ATB1071 treatment reduced infarct volume by ∼67% in EBP1-reintroduced mice, compared with ∼52% in mock-expressing *Ebp1* CKO mice (Fig. 7d, e).

**Figure 7.**
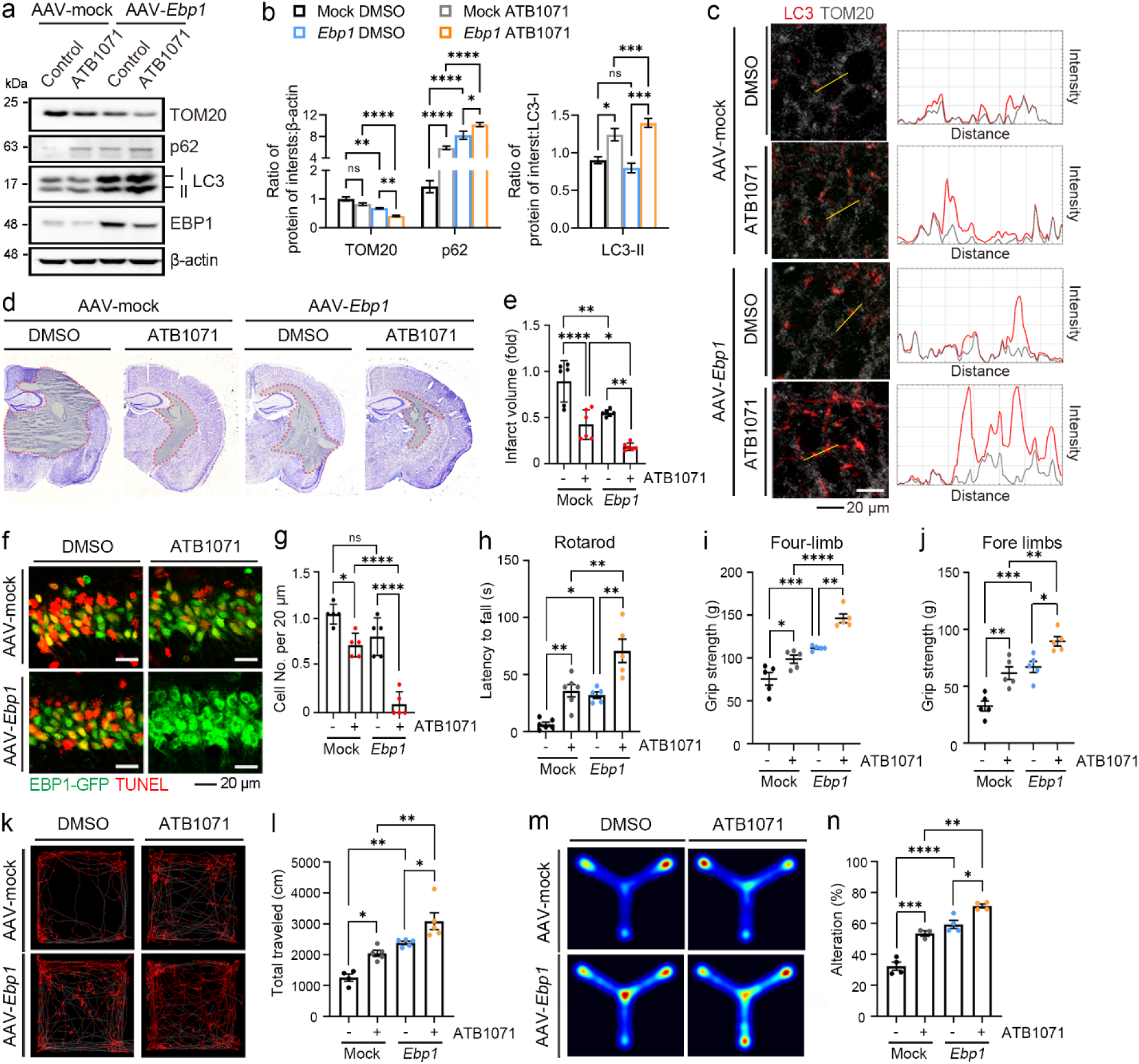
EBP1 reintroduction restores ATB1071-mediated mitophagy and neuroprotection after ischemic injury. **a** Immunoblotting analysis of hippocampal lysates of MCAO mice using the indicated antibodies. **b** Quantification of (**a**) (*n* = 4 mice). **c** Immunostaining of hippocampal sections from MCAO mice. The fluorescence intensity profiles along the yellow lines are displayed on the right. Scale bar, 20 µm. **d** Nissl staining of brain sections from the indicated groups. **e** Quantification of (**d**) (*n* = 6 mice). **f** TUNEL staining of the hippocampal CA1 region in GFP-expressing MCAO mice. **g** Quantification of (**f**) (*n* = 5 mice). **h-l** Motor-related behavioral assessments, including rotarod performance (**h**), four-limb grip strength (**i**), forelimb grip strength (**j**), and open-field locomotor activity (**k**) and (**l**) (n = 4–6 mice). **m**, **n** Y-maze spontaneous alternation test. The percentage of 1-2-3 order alterations was analyzed (*n* = 4 mice). For (**h**), (**i**) and (**j**), median values of triplicate trials per mouse are shown. Data are presented as mean ± SEM. Statistical significances were calculated using one-way ANOVA. \**p* < 0.05; \*\**p* < 0.01; \*\*\**p* < 0.001; \*\*\*\**p* < 0.0001; ns, not significant.

TUNEL staining further demonstrated that EBP1 reintroduction reduced neural cell death by ∼86% relative to the mock group following ATB1071 treatment (Fig. 7f, g). Assessment of neuromuscular function revealed that EBP1 reintroduction improved rotarod performance by ∼2.2-fold (Fig. 7h) and increased four-limb and forelimb grip strength by ∼1.5-fold in ATB1071-treated *Ebp1* CKO mice (Fig. 7i, j). Open-field and Y-maze tests additionally showed that EBP1-reintroduced mice treated with ATB1071 exhibited a ∼1.5-fold greater ambulatory activity and a ∼33% improvement in working memory compared with mock-expressing *Ebp1* CKO mice (Fig. 7k-n). These results demonstrate that EBP1 restores ATB1071-mediated mitophagy and enhances structural and functional recovery after cerebral IR injury.

### PK and toxicological evaluation support preclinical development of ATB1071

To evaluate the suitability of ATB1071 as a preclinical therapeutic for mitochondrial dysfunction, we characterized its PK properties. At an oral dose of 10 mg/kg, ATB1071 achieved a systemic exposure (AUC_last_) of 5,750.6 ng·h/mL, a terminal half-life (T /) of 9.8 hours, and an oral bioavailability of 70.4% (Supplementary Fig. 10a). The compound displayed efficient central nervous system (CNS) penetration, with a brain-to-plasma (B/P) ratio of 2.12 corresponding to a BBB penetration of 67.9% (Supplementary Fig. 10b, c). In both mice and dogs, ATB1071 exhibited dose-normalized exposures indicative of favorable bioavailability and moderate clearance. Toxicological assessments further supported a broad safety margin. In ICR mice, daily oral dosing for one week resulted in a no-observed-adverse-effect level (NOAEL) of at least 30 mg/kg. In rats, a two-week, three-times-per-week (TIW) regimen established a NOAEL of at least 300 mg/kg (Supplementary Fig. 10d). A cardiovascular telemetry study achieved a NOAEL of ∼100 mg/kg (Supplementary Fig. 10e). Genotoxicity studies reinforced the benign safety profile. An AMES assay across four *Salmonella* strains and one *Escherichia coli* strain showed no mutagenic activity (Supplementary Fig. 10f), and a chromosomal aberration assay detected no clastogenic effects. These results demonstrate that ATB1071 achieves robust systemic and CNS exposure, exhibits consistent PK behavior across species, and possesses a wide therapeutic window without detectable genotoxic or cardiac liabilities. Because ATB1071 induces mitophagy at concentrations of 2–3 μM (Fig. 2i, j, Supplementary Fig. 5h), which are attainable in the brain following an oral dose of 3 mg/kg (Supplementary Fig. 10a), these PK and safety data support the preclinical development of ATB1071 for diseases associated with mitochondrial dysfunction.

## DISCUSSION

This study defines a chemical N-degron strategy for activating p62-mediated mitophagy. Pharmacological enhancement of mitophagy has emerged as a promising therapeutic approach for a wide spectrum of diseases, including neurodegenerative disorders, metabolic syndromes, cardiovascular diseases, muscular dystrophies, and age-related pathologies. Despite decades of effort, however, most mitophagy inducers developed to date act by directly or indirectly modulating the mTOR and/or AMPK pathway rather than by engaging mechanisms that truly discriminate damaged mitochondria. Moreover, many of these compounds fail to achieve selective targeting of dysfunctional organelles, raising concerns about off-target effects and unintended metabolic consequences. Our findings address this gap by showing that p62 can be pharmacologically activated through N-degron mimicry to promote selective mitochondrial clearance under stress conditions.

In this study, we demonstrate that p62 mediates mitophagy as an N-recognin whose activity is allosterically enhanced by Arg/N-degrons. Building on this biochemical principle, we developed a mitophagy inducer that engages the p62-ZZ domain and activates p62 on damaged mitochondria without interfering with mTOR-dependent bulk autophagy. ATB1071 was designed to engage the p62-ZZ domain through N-degron interactions, thereby promoting PB1-driven self-polymerization and LIR-mediated recruitment of LC3-positive phagophores. During this N-degron–based process, ATB1071-induced p62 oligomers are selectively recruited to damaged mitochondria, in part via interactions with NIPSNAP1 and NIPSNAP2 that are exposed on the mitochondrial surface. These interactions appear to be at least partially independent of Parkin-mediated ubiquitination and instead rely on multivalent contacts among oligomeric species. Thus, ATB1071 reveals that chemical N-degrons can activate an endogenous autophagy receptor to couple damaged mitochondria with the autophagy machinery.

In mitophagy, p62 can bind ubiquitinated mitochondrial proteins and tether them to LC3 on autophagosomes to aid cargo recruitment^8, 63^. Despite its biochemical role, however, p62 has generally been viewed as dispensable for PINK1-Parkin–mediated mitophagy, largely because mitochondria can still be eliminated in its absence, and receptors such as NDP52 and OPTN perform more essential roles^13^. The current study revisits this assumption in light of our previous finding that p62 functions as an N-recognin regulated by Arg/N-degrons^14, 15^. Reassessing p62 as an N-recognin, we found that p62 is, under our experimental conditions, required for optimal mitophagy in both Parkin-independent and Parkin-dependent contexts. Moreover, we show that ATE1-dependent generation of Arg/N-degrons is essential for p62-mediated mitochondrial clearance. These findings provided the rationale for developing ATB1071, a chemical Arg/N-degron, as a mitophagy inducer. These results reposition p62 from a supporting mitophagy adaptor to a regulatable autophagy receptor whose activity can be enhanced by physiological or chemical N-degrons.

Our data suggest a model in which N-degron activation enables p62 to assemble on damaged mitochondria through damage-exposed adaptor platforms. Previous work showed that Arg/N-degron binding to the p62-ZZ domain triggers a conformational change that exposes the PB1 domain, thereby promoting p62 self-polymerization and oligomer formation through liquid-liquid phase separation (LLPS)^15, 45, 46^. In this study, ATB1071 promoted p62 interactions with NIPSNAP1/2, which are exposed on damaged mitochondria, and these interactions required the ZZ and PB1 domains but not the UBA domain. This indicates that ATB1071-driven p62 recruitment to NIPSNAP1/2 is mediated primarily through the N-degron-sensitive self-assembly axis rather than canonical ubiquitin binding. In parallel, EBP1 provided a Parkin-dependent platform for ATB1071-induced mitophagy during cerebral IR injury. Together, NIPSNAP1/2 and EBP1 represent distinct damage-associated mitochondrial platforms that converge on activated p62 to drive selective mitophagy.

LS remains a devastating mitochondrial disorder with significant unmet medical needs, as there is currently no effective drug available. Because the condition stems from OXPHOS defects and profound mitochondrial dysfunction, existing supportive measures cannot adequately address the underlying bioenergetic failure. Studies have shown that rapamycin significantly enhances the survival and attenuates disease progression in *Ndufs4 ^/^* mice deficient in OXPHOS complex I by delaying neurological onset, reducing brain lesions and neuroinflammation^64, 65^. While these studies provide supporting evidence for autophagy-based therapeutic strategies, their clinical applicability is limited by the broad effects of mTOR inhibition on bulk autophagy, metabolism, and inflammation. In this study, we demonstrate that pharmacological modulation of p62 using 20 doses of 10 mg/kg ATB1071 accelerates mitophagy in the brains of *Ndufs4*^-/-^ mice. ATB1071 administration also ameliorated neuroinflammation in *Ndufs4*^-/-^ brains. as shown by reduced GFAP^+^ astrocytes and Iba1^+^ microglia, and further supported by transcriptome sequencing. These therapeutic benefits were associated with marked improvements in survival and lifespan as well as muscle strength, neuromuscular coordination, and skin inflammation-associated hair loss. These results demonstrate that ATB1071, an mTOR-independent mitophagy inducer, ameliorates neuronal damages and disease-associated inflammatory phenotypes in *Ndufs4 ^/^* mice.

Cerebral IR injury remains a major clinical challenge, as despite advances in thrombectomy and thrombolysis, no approved treatment directly targets the mitochondrial dysfunction and neuronal death that drive long-term disability after stroke. We therefore employed ATB1071 as a therapeutic strategy to enhance mitochondrial quality control during the reperfusion phase. Our findings demonstrate that ATB1071 provides neuroprotection against IR injury by activating mitophagy. ATB1071 administration to MCAO-subjected mice markedly increased biochemical and cellular indicators of mitophagy, including reduced TOM20 abundance, elevated LC3 and p62 levels, and enhanced recruitment of LC3^+^ phagophores to damaged mitochondria. This mitophagy activation was accompanied by substantial neuroprotective outcomes, as evidenced by reduced brain infarct volume by ∼85% and neuronal death in the CA1 region by ∼91%, together with improvements in multiple behavioral and functional parameters affected by IR injury. Specifically, ATB1071 increased rotarod performance nearly fourfold, enhanced both four-limb and forelimb grip strengths, improved locomotor activity and center exploration in the open-field test, and restored working memory performance in the Y-maze. These results demonstrate that ATB1071 markedly attenuates tissue damage and restores neurological function following cerebral IR injury, supporting mitophagy activation as a neuroprotective strategy during reperfusion-associated mitochondrial stress. Although future studies will be needed to define the optimal treatment window and dosing regimen, these findings suggest that pharmacological activation of p62-mediated mitophagy may provide a therapeutic approach for ischemia-reperfusion injury.

EBP1 has been shown to be a multifunctional protein involved in various cellular processes including neuronal survival, differentiation, and brain development^67, 68^. More recently, the neuroprotective roles of EBP1 have been highlighted, as exemplified by studies showing that loss of *Ebp1* contributes to progression of sporadic AD due to amyloid β deposition and exacerbates neuronal death under IR stress^8, 69^. Of note, EBP1 plays a pivotal role in the clearance of dysfunctional mitochondria via its ubiquitination by Parkin, thus alleviating IR injury^8^. Our results highlight EBP1 as a key upstream regulator of ATB1071-induced p62-mediated mitophagy during cerebral IR injury. The loss of *Ebp1* not only abolished p62-mediated mitophagy but also sharply reduced the therapeutic benefits of ATB1071, resulting in limited suppression of infarct formation, diminished neuronal preservation, and poor recovery of motor, cognitive, and exploratory behaviors. Conversely, restoring EBP1 expression in *Ebp1* CKO brains restored p62-mediated mitophagy in response to ATB1071 and substantially enhanced structural and functional rescue. These findings identify EBP1 as a damage-associated mitochondrial platform that enables ATB1071-activated p62 to drive mitophagy and neuroprotection following cerebral IR injury.

Our preclinical efficacy studies in animal models of LS and cerebral IR injury raise the question of whether ATB1071 has a development path as an oral mitophagy activator. PK analysis shows that oral administration of ATB1071 at 3 mg/kg achieves a brain exposure of ∼2.7 μM, a concentration sufficient to induce mitophagy. Toxicology studies further demonstrate that the NOAEL is at least 300 mg/kg in rat following oral TIW dosing for two weeks, providing a substantial safety margin of ∼100-fold. Although longer and more frequent dosing regimens will likely be necessary in human clinical studies, our combined efficacy, PK, and toxicity findings support the view that ATB1071 is well positioned for further preclinical development in mitochondria-associated neurological diseases.

## Methods

### Ethics statement

All animal studies were conducted in accordance with guidelines set forth by the Institutional Animal Care and Use Committee (IACUC), Seoul National University (SNU-221202-1-5) and Sungkyunkwan University (SKKU-IACUC-2025-06-21-1).

### Reagents, compounds, and omics data analyses

The list of antibodies, reagents, and plasmids; the chemical synthesis and analytical data of ATB1071; and the mass spectrometric and transcriptomic analyses are described in the Supplementary Methods.

### Cells and cell culture

HeLa and SH-SY5Y cell lines were acquired from Korean Cell Line Bank. HEK293T cells were purchased from American Type Culture Collection (ATCC). *ATE1*^-/-^ and *p62*^-/-^ HeLa cells were gifted by Korea Research Institute of Bioscience and Biotechnology (KRIBB) and Dr. Eun-Kyeong Jo (Chungnam National University, Korea), respectively. Cell lines used in this study were cultured in Dulbecco’s Modified Eagle Medium (DMEM; Gibco, 11995073) supplemented with 10% fetal bovine serum (FBS; Gibco, 12483020) and maintained at 37°C and 5% CO2 in an incubator. Cells were routinely tested for mycoplasma contamination.

### Mice

The *Ndufs4*^+/-^ mice were obtained from the Jackson Laboratory (027058). *Ndufs4*^-/-^ mice were produced from their heterozygous parents and weaned at ∼21 days of age. *Ndufs4*^+/-^ and WT mice were pooled as controls for the experiments throughout the study, as they are identical in previous reports. The mouse *Pa2g4* gene is located on chromosome 10 (NM_011119). *pa2g4*-knockout mice were generated in collaboration with genOway (Lyon, France). To achieve neuron-specific deletion of *pa2g4*, homozygous mutants of the *pa2g4* allele (*pa2g4*^flox/flox^) were crossed with *CamK*Ⅱ*-Cre* mice ^8^. All mice were maintained under specific pathogen-free conditions. All mice employed for the experiments were weighed daily, and the mice that showed a 20% drop in maximum body weight were euthanized.

### Mitophagy assay

Cultured mt-mKeima HeLa cells were collected by trypsinization, and then resuspended in phenol red-free DMEM (Gibco, 21063029) supplemented with 10% FBS. The cells were centrifuged at 1,000 rpm for 2 min and resuspended in HBSS (Gibco, 14025092) supplemented with HEPES (Gibco, 15630080), followed by flow cytometer analysis. Lysosomal mt-mKeima was monitored by dual excitation ratiometric pH measurements using 488 nm (pH 7) and 561 nm (pH 4) lasers with 670/30 nm and 610/20 nm detection filters, respectively. Data were analyzed using FlowJo v 10.9.0. For confocal imaging, mt-mKeima HeLa cells were cultured in a 6-well glass-bottom plate. After mitophagy induction, the plate wells were washed with PBS, and filled with FluoroBrite DMEM (Gibco, A1896701) supplemented with HEPES. Images were acquired using an LSM 700 (Zeiss) with the same detection filter above. To visualize mitophagy induction in primary neurons, the mitophagy detection kit according to the manufacturer’s protocol.

Lysosomal mitophagy dye (Dojindo, MD01-10) was visualized using 561 nm lasers with 695/40 nm detection filter.

### Middle cerebral artery occlusion (MCAO)

Control (*Pa2g4*^flox/flox^) and CKO (*CamK*Ⅱ*-Cre*; *Pa2g4*^flox/flox^) mice aged 6–8 weeks were induced to cerebral ischemia injury using the transient middle cerebral artery occlusion (MCAO) operation. Mice were anesthetized with 4% isoflurane for induction, and the core body temperature was maintained at 37°C using a feedback-controlled heating system. Following a midline cervical incision, the common carotid artery (CCA) of the left side was carefully isolated and transiently occluded using microvascular clamps to induce forebrain ischemia for 30 minutes. Reperfusion was initiated by removing the clamps, and mice were monitored during the 72-hour post-reperfusion period. Sham-operated controls underwent identical surgical procedures without vascular occlusion. ATB1071 (10 mg/kg) was administered intraperitoneally at four time points post-reperfusion: immediately (0 h), 3 h, 1 day, and 2 days. After 72 h, mice were sacrificed and transcardially perfused with ice-cold PBS, followed by 4% PFA in PBS. For histological and biochemical analyses, brains were post-fixed in 4% PFA overnight at 4°C, followed in 30% sucrose (Duchefa, S0809) in PBS for 48 hours. Tissues were embedded in optimal cutting temperature compound (OCT; Sakura Finetek, 4583), snap-frozen on dry ice, and sectioned coronally at 20 μm using a Cryostat (Leica, CM1100).

### Generation of the mt-mKeima HeLa cell line

HEK293T cells for recombinant lentivirus production were seeded in a 100 mm^2^ dish before transfection. 6 μg psPax2 (Addgene, 12260), 2 μg pMD2-G (Addgene, 12259), and 8 μg pHAGE-mt-mkeima (Addgene, 131626) plasmid were incubated in Opti-MEM (Gibco, 31985070) with Lipofectamin 2000 (Thermo Fisher Scientific, 11668019). This transfection mix was added to the HEK293T cells. The virus was harvested and filtered with a polyethersulfone (PES) syringe filter, followed by polybrene (Sigma-Aldrich, TR-1003-G) supplements. HeLa cells were transduced with the virus mix. The fluorescent signal was monitored under a fluorescent microscope on Day 2 after infection. The mt-mKeima expressing HeLa cells were sorted using a flow cytometry-based cell sorter (Sony, SH800S).

### Primary neuron culture

Primary neurons were prepared from the brains of *Ndufs4*^-/-^ and control mouse pups. Genotyping was performed using Extract-N-Amp Tissue PCR Kit (Sigma-Aldrich, E3004) according to the manufacturer’s instructions. The genotyped pups were anesthetized and sacrificed by decapitation. The brains were dissected on ice to isolate the cortex, while the cerebellum, pons, and brain stem were removed. Neurons were dissociated by trypsinization and plated onto poly-d-lysine-coated either 6-well glass-bottom plates or 12-well plates. Primary neurons were maintained in Neurobasal A medium (Gibco, 21103-049) containing 2% B27 (Gibco, 17504-044), 0.5 mM glutamax (Gibco, 35050061). To eliminate glial cells, cultures were treated with 5-fluoro-2 deoxyuridine (Sigma-Aldrich, F0503) and Uridine (Sigma-Aldrich, U3003) at a final concentration of 20 μM as mitotic inhibitors. Fresh media containing the mitotic inhibitors were replaced every 3 to 4 days. Cells at 14 DIV were employed for the assays throughout the study.

### SDS-PAGE and immunoblotting

Cultured cells were washed twice with ice-cold PBS and lysed with RIPA buffer (50 mM Tris-HCl, 150 mM NaCl, 1% NP-40, 1% sodium deoxycholate and 0.1% SDS) (Biosesang, RC2002-050-00) supplemented with protease inhibitor (Abbkine, BMP1001) and phosphatase inhibitor cocktail (Sigma-Aldrich, P5726). Subsequently, the whole cell lysates were centrifuged at 13,000 rpm for 30 min at 4°C. The protein concentration in the supernatant was measured using the Bicinchoninic acid (BCA; Thermo Fisher Scientific, 23227) assay. 5X Laemmli sample buffer was added, followed by boiling for 10 min at 95°C. A same amount of proteins was loaded on the polyacrylamide gel and separated by sodium dodecyl sulfate (SDS)-electrophoresis. The proteins were transferred onto the polyvinylidene difluoride (PVDF; Millipore, IPVH000) membranes. The blots were blocked with 5% nonfat dry milk or BSA in PBS-T solution (20 mM Tris-HCl, pH 7.5, 150 mM NaCl, and 0.05% Tween 20). The blots were incubated with primary antibodies, followed by incubation with horseradish peroxidase-conjugated secondary antibodies. Immunoreactive bands were visualized by using SuperSignal West PICO PLUS (Thermo Fisher Scientific, 32106).

### Transfection and RNA interference

Plasmids were transfected into the cells by using Lipofectamine 2000 (Thermo Fisher Scientific, 11668019), and siRNAs were transfected at a final concentration of 80 nM using Lipofectamine RNAiMAX (Thermo Fisher Scientific, 13778075). The lipofectamine reagents with plasmids or siRNAs were incubated in a separate microtube with Opti-MEM (Gibco, 31985070) for 5 min, followed by mixing and incubation for 15 min. The transfection mix was added to the cells.

### In vitro p62 oligomerization assay

Cells were resuspended with lysis buffer (50 mM HEPES pH 7.4, 0.15 M KCl, 0.1% Nonidet P-40, 10% glycerol) supplemented with protease inhibitor cocktail. The cells were lysed by 10 cycles of freezing and thawing, followed by centrifugation at 13,000 rpm for 20 min at 4°C. The protein concentration in the supernatant was determined using the BCA assay. 5 μg of protein was incubated with 1 mM of ATB1071 in the presence of 100 μM bestatin (Thermo Fisher Scientific, 78433) at room temperature for 2 h. After incubation, Non-reducing 4X LDS sample buffer was added to the sample lysate, followed by boiling for 10 min at 95°C. The samples were loaded on a 3% stacking and 8% separating SDS-PAGE.

### RT-qPCR

Cells were lysed with 500 μl TRIzol (Invitrogen, 15596026) at room temperature for 5 min. Chloroform was added to the lysate and gently mixed, followed by centrifugation at 13,000 rpm for 15 min at 4°C. After transferring the supernatant to new tubes, same volume of isopropanol was added and mixed vigorously. The mixture was centrifuged at 13,000 rpm for 15 min at 4°C, and the supernatant was removed. 75% ethanol was added to the pellets and then centrifuged 13,000 rpm for 5 min at 4°C. The pellets were dried at room temperature and dissolved within RNase-free water. PrimeScript™ 1st strand cDNA Synthesis Kit (Takara, 6110A) was employed for the cDNA synthesis according to the manufacturer’s instruction. After dilution of the synthesized cDNA in ultrapure water at 1:7 ratio, qPCR master mix containing SYBR green (Smobio, TQ1210) and the following primers were mixed: *GAPDH* (5′-TCAACAGCGACACCCACTCC-3’; reverse, 5′-TGAGGTCCACCACCCTGTTG-3′), *PLSCR3* (forward, 5’-ACCCGGGCTCTTCTGGG-3’; reverse, 5’- GGCTGGGGTTTTCGAGATGA-3’), *NIPSNAP1* (forward, 5’-CTCGCGCTGGGGACGTT-3’; reverse, 5’-GTAGGCATCCAGGTATTCAGGC-3’), *NIPSNAP2* (forward, 5’-GCTCCGGACATGGACATCTT-3’; reverse, 5’-TGCTTCTAGGCATTCCGGTT-3’), *RBX1* (forward, 5’- TGTTTCCAAAATGGCGGCAG-3’; reverse, 5’-TTCCTGCAGATGGCACAGTT-3’).

For C*LS*, AccuTarget™ Human Real-Time PCR Primer (Bioneer, PHS-P01) was used. qPCR was performed using a BIO-RAD CFX Connect.

### Co-immunoprecipitation

Cells were transfected with the indicated plasmid using Lipofectamine 2000. The cells were collected and lysed with immunoprecipitation buffer (50 mM Tris-HCl pH 7.5, 150 mM NaCl, 0.5% Triton X-100, 1 mM EDTA) supplemented with 1 mM phenylmethylsulfonyl fluoride (PMSF; Thermo Fisher Scientific, 36978) and protease inhibitor cocktail. The cells were centrifuged at 13,000 rpm for 20 min. The protein concentration in the supernatant was measured using the BCA assay. The lysates were incubated with primary antibodies or Anti-c-Myc Agarose beads (Thermo Fisher Scientific, 20168) on the rotator overnight at 4°C. Protein A/G agarose beads were added to the lysates with primary antibodies and incubated on the rotator for 2 h. The beads were centrifuged at 4000 rpm for 1 min, and the supernatant was removed. The beads were subsequently washed 3 times with the lysis buffer. The isolated proteins were eluted with the 2X Laemmli sample buffer by boiling for 10 min at 95°C. The eluted proteins were subjected to immunoblotting analysis describe above.

### Cell viability assay

A water-soluble tetrazolium salt (WST)-based EZ-Cytox cell viability assay kit (Dogen, EZ-1000) was employed for the cell viability. HeLa and SH-SY5Y cells in a 96-well plate were treated with the YTK2205 or ATB1071 at the indicated concentration. Subsequently, assay reagent solution was added to each well using a multi-channel pipette. The cells were incubated for 2 h at 37°C in a CO_2_ incubator. Optical density (OD) values were measured at 450 nm using a microplate reader (Tecan, AT/Sunrise TW).

### Brain tissue preparation

Mice were anesthetized, and then perfused with PBS. Brains were rapidly extracted and fixed with 4% paraformaldehyde (PFA; Biosesang, PC2031-050-00) for 48 h at 4°C. After washing 3 times with PBS, the brains were sectioned sagittally or coronally into 30 μm thickness using a vibratome (Leica, VT1000S). The brain sections were transferred to brain tissue storage buffer (25 mM NaH_2_PO_4_ · 2H_2_O, 20 mM NaOH, 30% glycerol, and 30% Ethylene glycol) as a preservative and stored at -20°C.

### Immunofluorescence

Cells were cultured on coverslips in 12-well plates. Cells were fixed with 4% PFA for 15 min at room temperature, followed by permeabilization with 0.1% Triton X-100 (Biosesang, TR1020-500-00) buffer for 15 min. The cells were blocked with 2% BSA in PBS solution for 1 h at room temperature. Subsequently, the cells were incubated with primary antibodies diluted in the blocking solution overnight at 4°C. After washing 3 times with PBS, the cells were incubated with Alexa Fluor-conjugated secondary antibody diluted in the blocking solution for 1 h at room temperature. Using a DAPI-containing mounting solution (Vector Laboratories, H-1500-10), the coverslips were mounted on glass slides. Confocal images were acquired using the LSM 700 and analyzed with the Zeiss ZEN microscope software (blue edition). Brain sections were blocked with a PBS-based blocking solution (2% NGS, 1% BSA, 0.1 % Trition X-100, and 0.005 % Tween-20), followed by incubation with primary antibodies diluted in the blocking solution overnight at 4°C. After washing 3 times with PBS, the brain sections were incubated with the secondary antibody diluted in the blocking solution for 1 h. Subsequently, cover glasses were mounted on the glass slides using the DAPI-containing mounting solution. Images were acquired using an Axio scan7 (Zeiss).

### Nissl staining

Paraffin-embedded sections were deparaffinized twice in xylene for 10 min each, followed by gradual rehydration in decreasing concentrations of ethanol (100%, 95%, 90%, 70%, and 50%, 3 min each). After washing with PBS, sections were incubated in 0.25% Cresyl Violet acetate for 10 min. For gradual dehydration, sections were immersed in a series of increasing concentrations of ethanol (50%, 70%, 90%, 95%, and 100% ethanol, 3 min each) and xylene. The sections were mounted on slides using Permount (Fisher Chemical, SP15-100). Images were acquired using a slide scanner and analyzed with an Aperio Imagescope (Leica)

### Fluorescent dye staining

Cells were cultured in 6-well glass-bottom plates for imaging. The cells were incubated in HBSS containing 40 μM 2’,7’-dichlorofluorescin diacetate (DCFDA; Sigma-Aldric, D6883) at 37°C. The plate wells were washed with PBS and filled with FluoroBrite DMEM supplemented with HEPES. For quantitative analysis using the flow cytometer, SH-SY5Y cells were collected by trypsinization, and then resuspended in phenol red-free DMEM supplemented with 10% FBS. The cells were centrifuged at 1,000 rpm for 2 min, followed by resuspension in HBSS containing 100 nM tetramethylrhodamine ethyl ester perchlorate (TMRE; Med Chem Express, HY-D0985A) or 40 μM DCFDA, followed by incubation at 37°C. After washing with PBS, the cells were analyzed using a flow cytometer. The fluorescence was monitored using 561 nm lasers with 586/14 nm or 488 nm lasers with 530/43 detection filters. Data were analyzed using FlowJo v 10.9.0.

### Subcellular fractionation

Subcellular fractionation was performed with either Mitochondria/Cytosol fractionation Kit (Abcam, ab65320). In brief, HeLa or SH-SY5Y cells were resuspended with 1X Cytosol Extraction Buffer Mix containing protease inhibitor, followed by incubation on ice for 10 min. The cells were homogenized and centrifuged at 1000 x g for 10 min, again the supernatant was centrifuged at 10.000 x g for 30 min. Subsequently, the pellet was saved as the mitochondrial fraction, and the supernatant was collected as the cytosol fraction. For mitochondrial fractionation in the brain, Mitochondria Isolation Kit for Tissue (Thermo Fisher Scientific, 89801) was used according to the manufacturer’s instruction. Brains were homogenized in Reagent A and then mixed with Reagent C, followed by centrifugation at 1000 x g for 10 min. The supernatant was centrifuged at 10,000 x g. The pellet was saved as the mitochondrial fraction, and the supernatant was collected as the cytosol fraction.

### Sodium carbonate extraction and Proteinase K protection assay

Isolated mitochondrial fraction was resuspended in PBS or 0.1 M Na_2_CO_3_ (pH 11.5) and incubated on ice for 30 min, followed by centrifugation at 16,000 x g for 15 min to generate “Supernatant” and “Pellet” fractions. For proteinase K protection assay, half the volume of each fraction was incubated with 15 μg/mL Proteinase K for 15 min on ice. Subsequently, the reaction was stopped by addition of 1 mM PMSF. The fractions were subjected to immunoblotting analysis.

### Proximity ligation assay

A rabbit anti-TOMM20 and a rabbit anti-LC3 antibody were conjugated with PLA oligonucleotides using Duolink In Situ Probemaker PLUS (Sigma-Aldrich, DUO92009) and MINUS (Sigma-Aldrich, DUO92010), respectively, according to the manufacturer’s protocol. Brain sections were permeabilized with a PBS-based buffer (2% NGS, 1% BSA, 0.1 % Trition X-100, and 0.005 % Tween-20), followed by incubation with Duolink blocking solution for 1 h at 37°C. The sections were incubated with the PLA oligonucleotides conjugated antibodies overnight at 4°C. After washing with PLA washing buffer, the sections were hybridized, ligated, and amplified by rolling circle. Cover glasses were mounted on the glass slides using the DAPI-containing mounting solution. Images were acquired using an Axio scan7 and analyzed with the Zeiss ZEN microscope software (blue edition).

### Bioenergetic analysis

Mitochondrial respiration was monitored using a Seahorse XFe24 analyzer according to the manufacturer’s instruction. The oxygen consumption rate (OCR) was measured under basal conditions, followed by the sequential injection of 1.5 μM oligomycin, 1 μM carbonyl cyanide p-trifluoromethoxyphenylhydrazone (FCCP), and 0.5 μM rotenone/antimycin A. ATP production, Basal Respiration, and Maximum Respiration were analyzed using Wave 2.6.4. ATP levels were measured using an ATP Assay Kit (Abcam, 83355) according to the manufacturer’s instructions.

### Behavior test

The rotarod test was performed to monitor motor coordination. Mice were placed on a rotating bar that accelerated from 3 rpm to 40 rpm (0.1 rpm/s). The latency time to fall from the rod was measured. The mice underwent two consecutive practice sessions. The test was performed thrice on the testing day, and the maximum times were calculated. The grip strength test was performed thrice on the testing day, and the mean of the trials was calculated. The open field test was used to assess exploratory and anxiety behaviors. The open field area (44.5 x 44.5 cm) was separated into the center (28.5 x 28.5 cm) and the periphery. The movements of the mice were recorded for 20 min and analyzed using the video tracking software EthoVision XT14 (Noldus). In the Y-maze test, a symmetrical maze was used. The exploring movements of mice were recorded for 10 min and analyzed using EthoVision XT14.

### Statistical analysis

Data are presented as mean ± SD or SEM from at least three independent experiments. Quantification of the immunoreactive bands was performed using ImageJ. Arbitrary unit (A.U.) of fluorescence was calculated based on the intensity value obtained from Zeiss ZEN microscope software (blue edition). Statistical significance calculation was performed with Graphpad Prism v 8.2.1. Differences with P < 0.05 were considered statistically significant (****P < 0.0001; ***P < 0.001; **P < 0.01; *P < 0.05).

## Acknowledgements

We thank Dr. Eun-Kyeong Jo (Chungnam National University) for providing *p62^-/-^* HeLa cells. PK properties were conducted at Chaon (Seoul, South Korea) and Sundia (Shanghai, China), and toxicological evaluation were performed at Biotoxtech (Cheongju, South Korea). Schematic illustration of the autophagic N-degron pathway and the mt-mKeima-based analysis was created using BioRender. This work was supported by the National Research Foundation of Korea funded by the Ministry of Science and ICT (grants NRF-2020R1A5A1019023, NRF-2021R1A2B5B03002614, RS-2025-16652968 to Y.T.K.; RS-2026-25469096 to C.H.J.; RS-2023-00273887 to S.C.K.; NRF-2019R1A2C1089497 to Y.S.S.; RS-2024-00456173 to D.H.H.) and by the Ministry of Education (grants RS-2023-00249464 to C.H.J.; RS-2024-00461291 to C.H.J. and S.B.K.). Additional support was provided by the Korea Dementia Research Project through the Korea Dementia Research Center, funded by the Ministry of Health and Welfare and the Ministry of Science and ICT (grants HU21C0157 to J.Y.A.; RS-2024-00332875 to Y.H.S.; RS-2024-00447844 to C.H.J.). S.C.K. gratefully acknowledges support from the ASAN Foundation Biomedical Science Scholarship.

## Author Contributions

S.C.K and B.S.K contributed to the conception and experimental design, collection and assembly of data, data analysis and interpretation, manuscript writing; E.J.J, and D.-h.P. conducted animal experiments; D.H.H., H.K., G.E.L., and M.J.L performed IP-MS; Y.S.S. generated mt-mKeima HeLa; D.Y.P, J.H.L., and E.H.C contributed to immunoblotting analyses; A.J.H., and Y.H.S contributed to data analysis and interpretation; W.D.J., D.E.K., and S.B.K., contributed to structural characterization of ATB1071 ;C.H.J contributed to characterization of drug profiling and manuscript writing; Y.T.K., and J.Y.A. contributed to the data analysis and interpretation, manuscript writing, financial support, and final approval of the manuscript.

## Data availability

The mass spectrometry data have been deposited to the ProteomeXchange via the PRIDE database with the dataset identifier PXD077204. The RNA sequencing data generated in this study have been deposited in the National Center for Biotechnology Information (NCBI) Gene Expression Omnibus (GEO) data repository with the accession code GSE329028. Source data are provided with this paper.

## References

1. Spinelli, J.B. & Haigis, M.C. The multifaceted contributions of mitochondria to cellular metabolism. Nature Cell Biology 20, 745–754 (2018).

2. Wen, H. et al. Mitochondrial diseases: from molecular mechanisms to therapeutic advances. Signal Transduction and Targeted Therapy 10, 9 (2025).

3. Niyazov, D.M., Kahler, S.G. & Frye, R.E. Primary Mitochondrial Disease and Secondary Mitochondrial Dysfunction: Importance of Distinction for Diagnosis and Treatment. Molecular Syndromology 7, 122–137 (2016).

4. Zong, Y. et al. Mitochondrial dysfunction: mechanisms and advances in therapy. Signal Transduction and Targeted Therapy 9, 124 (2024).

5. Narendra, D.P. et al. PINK1 Is Selectively Stabilized on Impaired Mitochondria to Activate Parkin. PLOS Biology 8, e1000298 (2010).

6. Yamano, K. & Youle, R.J. PINK1 is degraded through the N-end rule pathway. Autophagy 9, 1758–1769 (2013).

7. Sarraf, S.A. et al. Landscape of the PARKIN-dependent ubiquitylome in response to mitochondrial depolarization. Nature 496, 372–376 (2013).

8. Hwang, I. et al. PA2G4/EBP1 ubiquitination by PRKN/PARKIN promotes mitophagy protecting neuron death in cerebral ischemia. Autophagy 20, 365–379 (2024).

9. Park, S.-W. et al. Differential roles of N- and C-terminal LIR motifs in the catalytic activity and membrane targeting of RavZ and ATG4B proteins. BMB Rep. 57, 497–502 (2024).

10. Klionsky, D.J. et al. Guidelines for the use and interpretation of assays for monitoring autophagy (4th edition)(1). Autophagy 17, 1–382 (2021).

11. Rogov, V., Dötsch, V., Johansen, T. & Kirkin, V. Interactions between Autophagy Receptors and Ubiquitin-like Proteins Form the Molecular Basis for Selective Autophagy. Molecular Cell 53, 167–178 (2014).

12. Matsumoto, G., Shimogori, T., Hattori, N. & Nukina, N. TBK1 controls autophagosomal engulfment of polyubiquitinated mitochondria through p62/SQSTM1 phosphorylation. Human Molecular Genetics 24, 4429–4442 (2015).

13. Lazarou, M. et al. The ubiquitin kinase PINK1 recruits autophagy receptors to induce mitophagy. Nature 524, 309–314 (2015).

14. Cha-Molstad, H. et al. Amino-terminal arginylation targets endoplasmic reticulum chaperone BiP for autophagy through p62 binding. Nature Cell Biology 17, 917–929 (2015).

15. Cha-Molstad, H. et al. p62/SQSTM1/Sequestosome-1 is an N-recognin of the N-end rule pathway which modulates autophagosome biogenesis. Nature Communications 8, 102 (2017).

16. Novak, I. et al. Nix is a selective autophagy receptor for mitochondrial clearance. EMBO reports 11, 45–51-51 (2010).

17. Liu, L. et al. Mitochondrial outer-membrane protein FUNDC1 mediates hypoxia-induced mitophagy in mammalian cells. Nature Cell Biology 14, 177–185 (2012).

18. Wei, Y., Chiang, W.-C., Sumpter, R., Jr., Mishra, P. & Levine, B. Prohibitin 2 Is an Inner Mitochondrial Membrane Mitophagy Receptor. Cell 168, 224–238.e210 (2017).

19. Princely Abudu, Y., et al. NIPSNAP1 and NIPSNAP2 Act as “Eat Me” Signals for Mitophagy. Developmental Cell 49, 509–525.e512 (2019).

20. Gorman, G.S., et al. Mitochondrial diseases. Nature Reviews Disease Primers 2, 16080 (2016).

21. Stenton, S.L. et al. Leigh Syndrome: A Study of 209 Patients at the Beijing Children’s Hospital. Annals of Neurology 91, 466–482 (2022).

22. Lake, N.J., Compton, A.G., Rahman, S. & Thorburn, D.R. Leigh syndrome: One disorder, more than 75 monogenic causes. Annals of Neurology 79, 190–203 (2016).

23. Rahman, S. et al. Leigh syndrome: Clinical features and biochemical and DNA abnormalities. Annals of Neurology 39, 343–351 (1996).

24. Choi, M.L. et al. Pathological structural conversion of α-synuclein at the mitochondria induces neuronal toxicity. Nature Neuroscience 25, 1134–1148 (2022).

25. Mahmud, S.A., Qureshi, M.A. & Pellegrino, M.W. On the offense and defense: mitochondrial recovery programs amidst targeted pathogenic assault. The FEBS Journal 289, 7014–7037 (2022).

26. Meyer, J.N. et al. Mitochondria as a Target of Environmental Toxicants. Toxicological Sciences 134, 1–17 (2013).

27. Zhang, G. et al. Mitochondrial DNA leakage: underlying mechanisms and therapeutic implications in neurological disorders. Journal of Neuroinflammation 22, 34 (2025).

28. Li, X.-y., et al. Global, regional, and national burden of ischemic stroke, 1990-2021: an analysis of data from the global burden of disease study 2021. eClinicalMedicine 75 (2024).

29. Donkor, E.S. Stroke in the 21st Century: A Snapshot of the Burden, Epidemiology, and Quality of Life. Stroke Research and Treatment 2018, 3238165 (2018).

30. Zhang, M. et al. Ischemia-reperfusion injury: molecular mechanisms and therapeutic targets. Signal Transduction and Targeted Therapy 9, 12 (2024).

31. Fogo, G.M. et al. Mitochondrial dynamics and quality control regulate proteostasis in neuronal ischemia-reperfusion. Autophagy 21, 1492–1506 (2025).

32. Sanderson, T.H., Reynolds, C.A., Kumar, R., Przyklenk, K. & Hüttemann, M. Molecular Mechanisms of Ischemia–Reperfusion Injury in Brain: Pivotal Role of the Mitochondrial Membrane Potential in Reactive Oxygen Species Generation. Molecular Neurobiology 47, 9–23 (2013).

33. Li, W. et al. Ischemia - Reperfusion injury: A roadmap to precision therapies. Molecular Aspects of Medicine 104, 101382 (2025).

34. Li, Q. et al. Rapamycin attenuates mitochondrial dysfunction via activation of mitophagy in experimental ischemic stroke. Biochemical and Biophysical Research Communications 444, 182–188 (2014).

35. Ryu, D. et al. Urolithin A induces mitophagy and prolongs lifespan in C. elegans and increases muscle function in rodents. Nature Medicine 22, 879–888 (2016).

36. Thomson, A.W., Turnquist, H.R. & Raimondi, G. Immunoregulatory functions of mTOR inhibition. Nature Reviews Immunology 9, 324–337 (2009).

37. Asrani, K. et al. mTORC1 feedback to AKT modulates lysosomal biogenesis through MiT/TFE regulation. The Journal of Clinical Investigation 129, 5584–5599 (2019).

38. Kezic, A., Popovic, L. & Lalic, K. mTOR Inhibitor Therapy and Metabolic Consequences: Where Do We Stand? Oxidative Medicine and Cellular Longevity 2018, 2640342 (2018).

39. Cen, X. et al. Pharmacological targeting of MCL-1 promotes mitophagy and improves disease pathologies in an Alzheimer’s disease mouse model. Nature Communications 11, 5731 (2020).

40. Tan, S. et al. Targeted clearance of mitochondria by an autophagy-tethering compound (ATTEC) and its potential therapeutic effects. Science Bulletin 68, 3013–3026 (2023).

41. Okarmus, J. et al. USP30 inhibition induces mitophagy and reduces oxidative stress in parkin-deficient human neurons. Cell Death & Disease 15, 52 (2024).

42. Antico, O., Thompson, P.W., Hertz, N.T., Muqit, M.M.K. & Parton, L.E. Targeting mitophagy in neurodegenerative diseases. Nature Reviews Drug Discovery 24, 276–299 (2025).

43. Rosencrans, W.M. et al. Putative PINK1/Parkin activators lower the threshold for mitophagy by sensitizing cells to mitochondrial stress. Science Advances 11, eady0240 (2025).

44. Sriram, S.M., Kim, B.Y. & Kwon, Y.T. The N-end rule pathway: emerging functions and molecular principles of substrate recognition. Nature Reviews Molecular Cell Biology 12, 735–747 (2011).

45. Yoo, Y.D. et al. N-terminal arginylation generates a bimodal degron that modulates autophagic proteolysis. Proceedings of the National Academy of Sciences 115, E2716–E2724 (2018).

46. Ji, C.H. et al. The N-Degron Pathway Mediates ER-phagy. Molecular Cell 75, 1058–1072.e1059 (2019).

47. Jung, C.H. et al. The N-degron pathway mediates the autophagic degradation of cytosolic mitochondrial DNA during sterile innate immune responses. Cell Reports 44, 115094 (2025).

48. Shim, S.M. et al. The Cys-N-degron pathway modulates pexophagy through the N-terminal oxidation and arginylation of ACAD10. Autophagy 19, 1642–1661 (2023).

49. Heo, A.J. et al. The N-terminal cysteine is a dual sensor of oxygen and oxidative stress. Proceedings of the National Academy of Sciences 118, e2107993118 (2021).

50. Kwon, Y.T., Kashina, A.S. & Varshavsky, A. Alternative Splicing Results in Differential Expression, Activity, and Localization of the Two Forms of Arginyl-tRNA-Protein Transferase, a Component of the N-End Rule Pathway. Molecular and Cellular Biology 19, 182–193 (1999).

51. Tasaki, T. et al. The Substrate Recognition Domains of the N-end Rule Pathway*. Journal of Biological Chemistry 284, 1884–1895 (2009).

52. Kwon, D.H. et al. Insights into degradation mechanism of N-end rule substrates by p62/SQSTM1 autophagy adapter. Nature Communications 9, 3291 (2018).

53. Turco, E. et al. FIP200 Claw Domain Binding to p62 Promotes Autophagosome Formation at Ubiquitin Condensates. Molecular Cell 74, 330–346.e311 (2019).

54. Lee, Y.J. et al. Chemical modulation of SQSTM1/p62-mediated xenophagy that targets a broad range of pathogenic bacteria. Autophagy 18, 2926–2945 (2022).

55. Jung, E.J. et al. The N-degron pathway mediates lipophagy: The chemical modulation of lipophagy in obesity and NAFLD. Metabolism 146, 155644 (2023).

56. Katayama, H., Kogure, T., Mizushima, N., Yoshimori, T. & Miyawaki, A. A Sensitive and Quantitative Technique for Detecting Autophagic Events Based on Lysosomal Delivery. Chemistry & Biology 18, 1042–1052 (2011).

57. Zhang, Y. et al. ZZ-dependent regulation of p62/SQSTM1 in autophagy. Nature Communications 9, 4373 (2018).

58. Ortigoza-Escobar, J.D. et al. Ndufs4 related Leigh syndrome: A case report and review of the literature. Mitochondrion 28, 73–78 (2016).

59. Shil, S.K. et al. Ndufs4 ablation decreases synaptophysin expression in hippocampus. Scientific Reports 11, 10969 (2021).

60. Yoon, J.-Y. et al. Metabolic rescue ameliorates mitochondrial encephalo-cardiomyopathy in murine and human iPSC models of Leigh syndrome. Clinical and Translational Medicine 12, e954 (2022).

61. Li, N. et al. Mitochondrial Complex I Inhibitor Rotenone Induces Apoptosis through Enhancing Mitochondrial Reactive Oxygen Species Production*. Journal of Biological Chemistry 278, 8516–8525 (2003).

62. Chu, C.T. et al. Cardiolipin externalization to the outer mitochondrial membrane acts as an elimination signal for mitophagy in neuronal cells. Nature Cell Biology 15, 1197–1205 (2013).

63. Yamada, T. et al. Mitochondrial Stasis Reveals p62-Mediated Ubiquitination in Parkin-Independent Mitophagy and Mitigates Nonalcoholic Fatty Liver Disease. Cell Metabolism 28, 588–604.e585 (2018).

64. Johnson, S.C. et al. mTOR Inhibition Alleviates Mitochondrial Disease in a Mouse Model of Leigh Syndrome. Science 342, 1524–1528 (2013).

65. Martin-Perez, M. et al. PKC downregulation upon rapamycin treatment attenuates mitochondrial disease. Nature Metabolism 2, 1472–1481 (2020).

66. Rho, H., Kim, U. & Song, J. Ubiquitination and ubiquitin-like modifications in metabolic dysfunction-associated steatotic liver disease: mechanisms and implications. BMB Rep. 58, 371–388 (2025).

67. Ko, H.R. et al. Neuron-specific expression of p48 Ebp1 during murine brain development and its contribution to CNS axon regeneration. BMB Rep 50, 126–131 (2017).

68. Ko, H.R. et al. Roles of ErbB3-binding protein 1 (EBP1) in embryonic development and gene-silencing control. Proc Natl Acad Sci U S A 116, 24852–24860 (2019).

69. Kim, B.-S. et al. EBP1 potentiates amyloid β pathology by regulating γ-secretase. Nature Aging 5, 486–503 (2025).

70. Yoon, J.G. et al. De novo missense variants in HDAC3 leading to epigenetic machinery dysfunction are associated with a variable neurodevelopmental disorder. The American Journal of Human Genetics 111, 1588–1604 (2024).

